# A Grad-seq view of RNA and protein complexes in *Pseudomonas aeruginosa* under standard and bacteriophage predation conditions

**DOI:** 10.1101/2020.12.06.403469

**Authors:** Milan Gerovac, Laura Wicke, Kotaro Chihara, Cornelius Schneider, Rob Lavigne, Jörg Vogel

## Abstract

The Gram-negative rod-shaped bacterium *Pseudomonas aeruginosa* is not only a major cause of nosocomial infections but also serves as a model species of bacterial RNA biology. While its transcriptome architecture and post-transcriptional regulation through the RNA-binding proteins Hfq, RsmA and RsmN have been studied in detail, global information about stable RNA–protein complexes is currently lacking in this human pathogen. Here, we implement Gradient profiling by sequencing (Grad-seq) in exponentially growing *P. aeruginosa* cells to comprehensively predict RNA and protein complexes, based on glycerol gradient sedimentation profiles of >73% of all transcripts and ∼40% of all proteins. As to benchmarking, our global profiles readily reported complexes of stable RNAs of *P. aeruginosa*, including 6S RNA with RNA polymerase and associated pRNAs. We observe specific clusters of non-coding RNAs, which correlate with Hfq and RsmA/N, and provide a first hint that *P. aeruginosa* expresses a ProQ-like FinO domain containing RNA-binding protein. To understand how biological stress may perturb cellular RNA/protein complexes, we performed Grad-seq after infection by the bacteriophage ΦKZ. This model phage, which has a well-defined transcription profile during host takeover, displayed efficient translational utilization of phage mRNAs and tRNAs, as evident from their increased co-sedimentation with ribosomal subunits. Additionally, Grad-seq experimentally determines previously overlooked phage-encoded non-coding RNAs. Taken together, the *Pseudomonas* protein and RNA complex data provided here will pave the way to a better understanding of RNA-protein interactions during viral predation of the bacterial cell.

**IMPORTANCE:** Stable complexes by cellular proteins and RNA molecules lie at the heart of gene regulation and physiology in any bacterium of interest. It is therefore crucial to globally determine these complexes in order to identify and characterize new molecular players and regulation mechanisms. Pseudomonads harbour some of the largest genomes known in bacteria, encoding ∼5,500 different proteins. Here, we provide a first glimpse on which proteins and cellular transcripts form stable complexes in the human pathogen *Pseudomonas aeruginosa*. We additionally performed this analysis with bacteria subjected to the important and frequently encountered biological stress of a bacteriophage infection. We identified several molecules with established roles in a variety of cellular pathways, which were affected by the phage and can now be explored for their role during phage infection. Most importantly, we observed strong co-localization of phage transcripts and host ribosomes, indicating the existence of specialized translation mechanisms during phage infection. All data are publicly available in an interactive and easy to use browser.

## INTRODUCTION

*Pseudomonas aeruginosa* is a Gram-negative environmental γ-proteobacterium and a critical life-threatening pathogen in humans with compromised immune defence (1). It is the main cause of death in cystic fibrosis patients (2) and hard to treat with antibiotics due to its diverse export capabilities and its ability to form strong biofilms (3). Unsurprisingly, the medical challenges associated with nosocomial infections and drug resistance of *P. aeruginosa* have prompted much effort to better understand the molecular biology of this important human pathogen. With a size of 6.3 Mbp and ∼5,570 predicted open reading frames (ORFs), the genome of *P. aeruginosa* is one of the largest among prokaryotes (4). These genes, many of which encode paralogous protein, endow the bacterium with a remarkable functional diversity that allows it to thrive in different habitats. Gene expression control also seems to be complex in *P. aeruginosa*: an unusually high 8.4% of all genes are predicted to encode regulatory proteins, foremost transcription factors.

In addition to extensive gene regulation at the DNA level, there has been increasing evidence for post-transcriptional regulation to play an important role in *P. aeruginosa* (5). The bacterium possesses homologs of two general RNA-binding proteins (RBPs), CsrA and Hfq, which work in conjunction with small regulatory RNAs (6). An important role of Hfq in *P. aeruginosa* was first predicted almost 15 years ago when an *hfq* knockout strain was observed to display altered levels for 5% of all transcripts (7). More recent work employing global RNA interactome techniques (8–11) suggests that Hfq interacts with a large number of coding and noncoding transcripts of *P. aeruginosa*. A distinct feature of Hfq-mediated regulation in *Pseudomonas* is that Hfq inhibits translation of target transcripts by forming a regulatory complex with the catabolite repression protein Crc (12, 13), likely by recognition of nascent transcripts (9).

The other major regulatory RBPs include the CsrA-like proteins RsmA and RsmN, which act as global translational repressors through recognizing GGA motifs in the 5’ region of mRNAs (14–18), partly so in a combinatorial fashion with Hfq (19). By contrast, much less is known about a putative homolog of FinO/ProQ-like proteins, which is an emerging new family of sRNA-related RBPs in Gram-negative bacteria (20–24). Judging by the presence of a FinO domain, *P. aeruginosa* does possess a candidate protein (25), but whether it is expressed and forms complexes with cellular transcripts remains unknown. Similarly, cold shock proteins (CSPs), which interact with hundreds if not thousands of transcripts in some bacteria (26), have not been studied in *P. aeruginosa*.

The past couple of years have witnessed the development of new methods to discover RBPs and study RNA-protein complexes and interactions on a global level (27), one of which is gradient profiling by sequencing, a.k.a. Grad-seq (22). Grad-seq partitions native cellular lysates, including RNA–protein complexes, according to their molecular weight and shape on a glycerol gradient. Subsequent fractionation and analysis by RNA-seq and mass spectrometry of each fraction enables visualization of in-gradient distributions of, ideally, all expressed RNAs and detectable soluble proteins from the growth condition of interest. The method has been successfully applied to several different bacteria, revealing in *Salmonella enterica* a previously unknown global activity of ProQ (22), new stable RNA and protein associations with the ribosome in *Escherichia coli* (28); and a mechanism of sRNA stabilization by 3’→5’ exonucleolytic trimming in the Gram-positive human pathogen *Streptococcus pneumoniae* (29). In all these studies, Grad-seq produced a previously unavailable resource for the comprehensive prediction of protein complexes, with or without RNA components, of a wide functional spectrum.

Despite its importance as a human pathogen and model bacterium, global data informing on RNA/protein complexes are lacking for *P. aeruginosa* as well as other pseudomonads. Here, we pioneer the application of bacterial complexomics in *P. aeruginosa*, analysing exponentially growing cultures of this bacterium with Grad-seq. Our analysis provides the first systems biology-based description of RNA-based housekeeping and regulatory systems in this species, which includes a stable RNA polymerase (RNAP)-6S RNA complex. We describe in-gradient distribution for thousands of mRNAs and sRNAs and predict hundreds of protein complexes involving metabolic and signal transduction proteins. In addition to performing Grad-seq on naïve bacteria, we apply the approach for the first time to visualize transcript associations of an invading phage (ΦKZ) and to identify phage-encoded regulatory RNAs. These data sets are provided along with an online browser that helps to visualize the obtained sedimentation profiles in a searchable fashion and allows for them to be compared with Grad-seq data from other Gram-negative and Gram-positive species.

## RESULTS

### Gradient fractionation of P. aeruginosa in early exponential growth phase

As a prerequisite for Grad-seq of *P. aeruginosa* (see workflow in **Fig. 1A**), we established a glycerol gradient analysis of RNA and proteins from a lysate of strain PAO1. We chose to analyse samples from the exponential phase of growth (OD_600_ of 0.3) in a rich culture medium to allow comparisons to previously established analyses under standard growth conditions. In addition, the exponential phase is optimal for phage infection and replication of ΦKZ (30). Phage predation represents a biotic stress for PAO1, relevant to the experimental design and narrative below. A cleared lysate was separated on a 10-40% glycerol gradient, followed by fractionation and gel-based analysis of either protein or RNA molecules in each of the 20 fractions obtained. In addition, the pellet was included in this analysis, as it contains translating ribosomes as well as aggregated and unfolded proteins.

**Figure 1.**
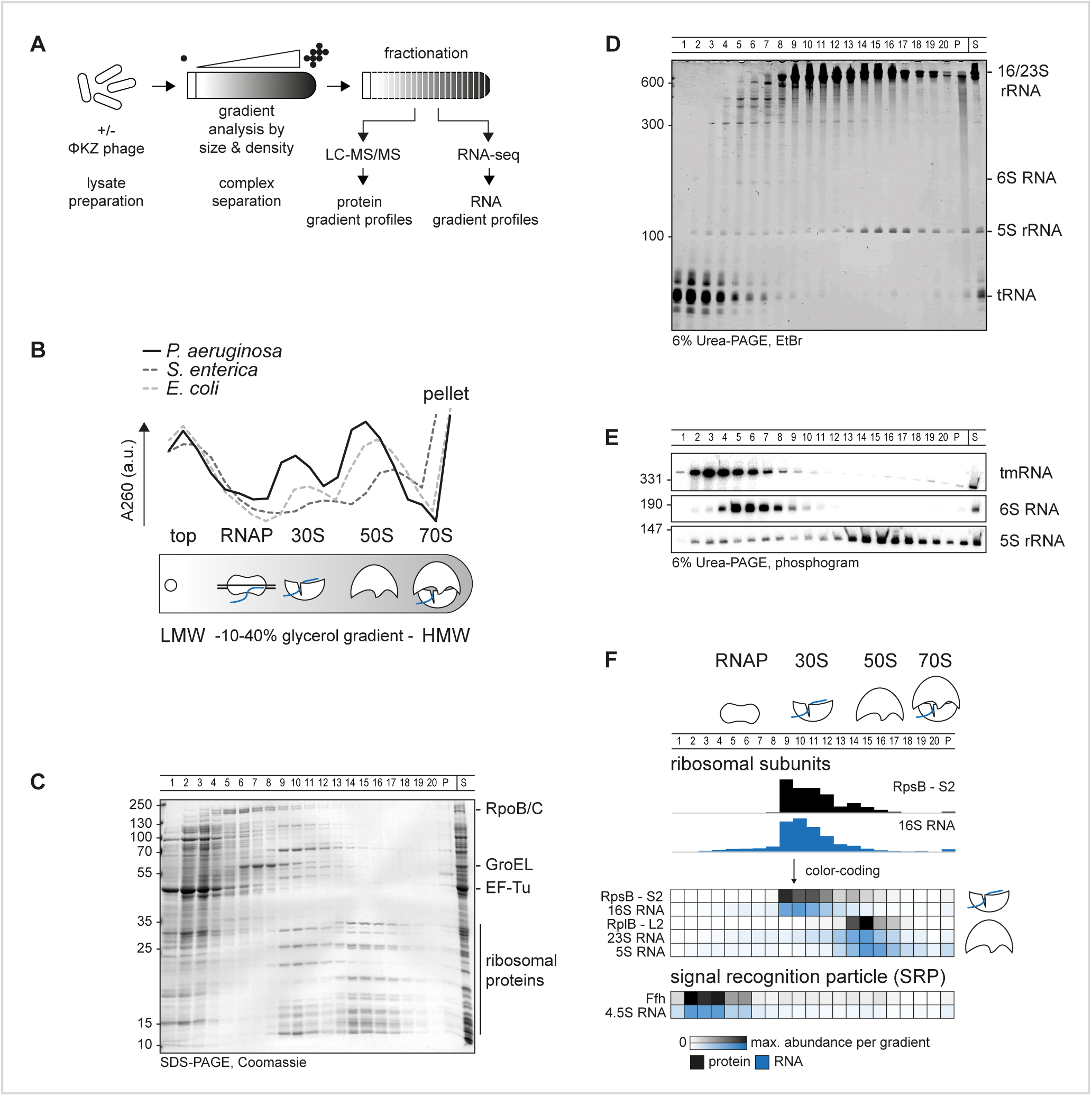
Grad-seq in *Pseudomonas*. **(A)** Cellular lysate is analysed in a gradient and fractionated. High-throughput analysis by mass-spectrometry (LC-MS/MS) and RNA-sequencing reveals sedimentation profiles. **(B)** Complexes are resolved in a 10-40% glycerol gradient by size and density. The 260 nm-absorption profile of *Pseudomonas* lysate indicates LMW fractions at the top, 30S small, 50S large ribosomal subunits at fractions 9 and 15, respectively, and an elevated absorption in the pellet at HMW fractions. The profile is shifted by about one fraction compared to *E. coli* and *S. enterica*. The RNAP sediments between the top and 30S subunit. **(C)** The apparent proteome resembles the positions in (B). EF-Tu is a transient translation factor and sedimented in the top, RNAP subunits RpoB/C and GroEL sediment before 30S fractions. **(D)** tRNAs sedimented in top fractions, ribosomal RNAs resemble well the distribution in (B,C), 6S RNA sedimentation correlated with RNAP in (C). **(E)** tmRNA sedimented in the pellet and at RNAP fractions, 6S RNA sedimentation correlated with RNAP fractions and 5S rRNA with 50S ribosomal subunit fractions. **(F)** High throughput determined sedimentation profiles matched well with in gel and previously published results in other species.

As a quality control, we determined the absorption profile of the gradient fractions at 260 nm (**Fig. 1B**). We observed the expected bulk peak at the top of the gradient, which corresponds to free proteins and RNAs in low-molecular-weight (LMW) fractions. Also, two pronounced peaks corresponding to the small (30S) and large (50S) ribosomal subunits in high-molecular-weight (HMW) fractions were observed. Compared to previously obtained profiles for Grad-seq of *E. coli* or *S. enterica* (22, 28), we observed much fewer 70S ribosomes. In addition, the ribosomal subunits were shifted by one fraction towards the top of the gradient, which might indicate species-specific disintegration of ribosomal subunit complexes, or a relaxed rRNA configuration in *Pseudomonas* in the present growth condition.

SDS-PAGE showed general distributions of abundant soluble proteins and their complexes within the gradient (**Fig. 1C**). As expected, translation factor EF-Tu is one of the most abundant proteins in *Pseudomonas* (**Table S1**). The EF-Tu protein, which transiently delivers transfer RNAs (tRNAs) to the ribosome (31), primarily sedimented in a ribosome-free manner, occurring in the LMW fractions 1-3. RNAP subunits were primarily found in fractions 5-7, whereas the major protein GroEL abounded in fractions 6-9 (i.e., complexes in the ∼400-800 kDa range). The 30S and 50S ribosomal subunits, with a molecular weight of ∼1 MDa and ∼2 MDa, respectively, sedimented in fractions 9-13 or 14-16.

Similarly, we stained abundant RNA molecules in the individual gradient fractions separated in a denaturing gel (**Fig. 1D**). As expected, tRNAs sedimented primarily in LMW fractions 1-4, whereas the abundant 16S and 23S/5S rRNA molecules coincided with the 30S or 50S proteins in fractions 9-13 or 14-17, respectively. As a benchmark, we also determined the sedimentation profiles of several stable RNAs by northern blot analysis (**Fig. 1E**). Transfer/messenger RNA (tmRNA, encoded by *ssrA*) forms a ribonucleoprotein particle (RNP) with the SmpB protein to rescue stalled ribosome by a trans-translation mechanism (32). Consistent with previous studies in *E. coli* and *S. enterica*, tmRNA sedimented in fractions 3-7 (22, 28). The 180-nt 6S RNA (33) accumulated in fractions 5-7, as expected from its reported association with RNAP in *E. coli* (34). In summary, these observations qualified the gradient samples for high-throughput analysis by Grad-seq, with the aim to obtain global sedimentation profiles of cellular RNA and protein molecules (**Fig. 1F**).

### Global predictions of macromolecular complexes for P. aeruginosa

For a global view of stable complexes in the exponential growth phase, we profiled the proteins and transcripts from each gradient fraction by mass spectrometry and RNA-seq, respectively (**Table S1**). Protein and RNA abundance was normalized using external spike-ins, enabling us to delimit the detection thresholds (**Fig. S1**). Targeted inspection of these high-throughput data readily showed expected co-occurrences of stable RNA species with their major protein partners (**Figs. 1F, S2**). For example, cDNA counts of the 16S/23S rRNAs peaked in the same fractions as did peptide counts of the respective ribosomal proteins, and 4.5S RNA of the signal recognition particle (SRP) peaked in the same fractions as did the major SRP protein Ffh. Another case in point is the 6S RNA-RNAP complex whose protein components (RpoA/B/C/Z encoding the α/β/β’/ω subunits of RNAP) aligned well with 6S RNA in gradient fractions 5-6 (**Fig. 2A**). Grad-seq even recovered the tiny pRNAs (35, 36) produced by RNAP as it disentangles itself from the 6S RNA (**Fig. 2B**). These established interactions provide additional confidence to the correlations described below. Mass spectrometry recovered 2,182 proteins in total, i.e., ∼40% of the annotated proteome of *P. aeruginosa* strain PAO1 (37), with these proteins showing a wide range of distribution in the gradient (**Figs. 3A, S2A**). While most of these proteins sedimented in LMW fractions 1-3, indicating that they are part of either no or only small complexes, there were also protein clusters in the HMW fractions, in addition to the expected two ribosomal subunit clusters. We will return to some of these HMW clusters in the next section.

**Figure 2.**
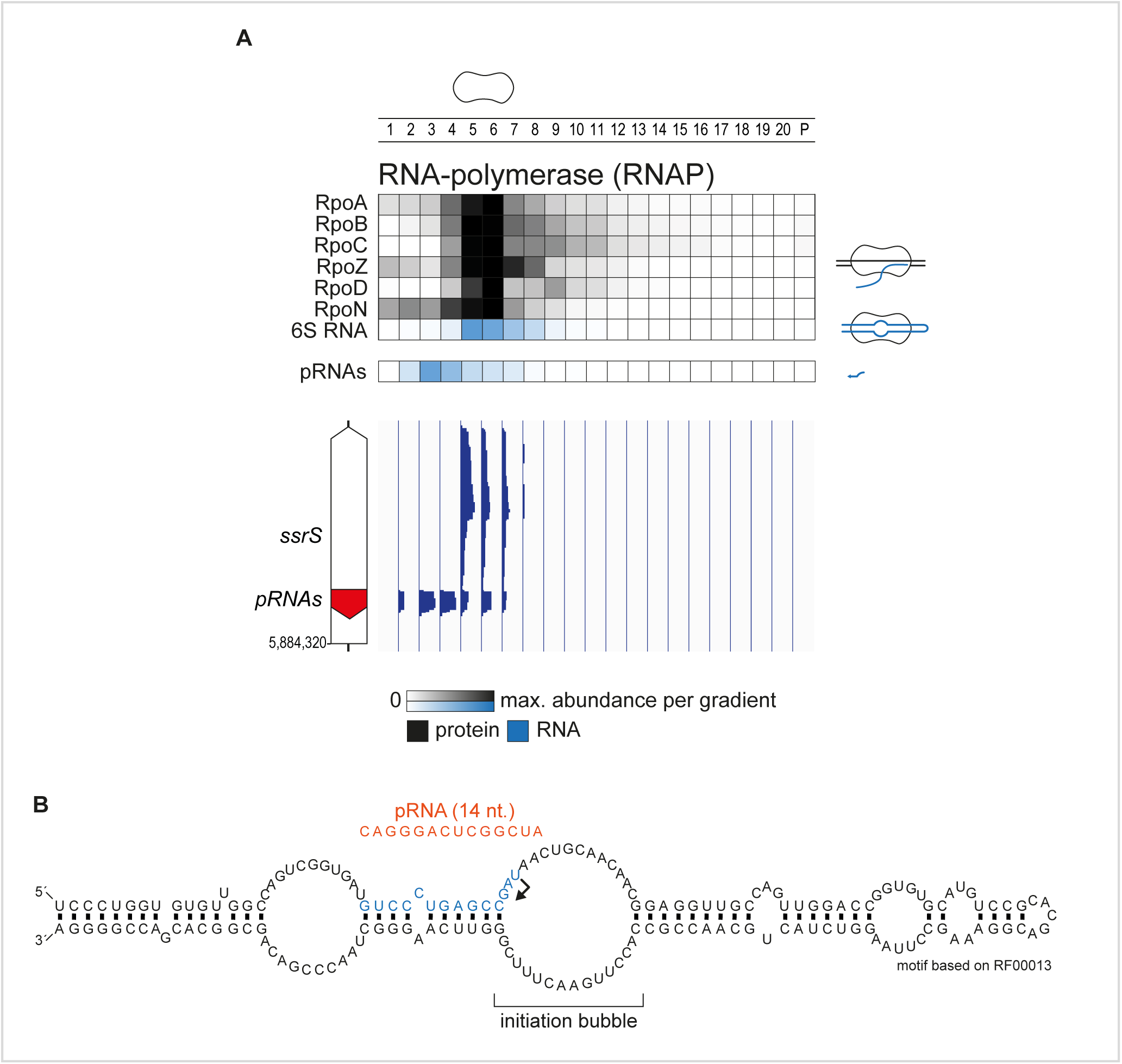
Grad-seq allows for mapping of pRNAs. **(A)** RNAP subunits, 6S RNA and pRNAs co-sedimented in the gradient. pRNAs were mapped by differential sedimentation. Importantly, pRNAs sedimented at a much larger position in fraction three indicating presence in a complex. **(B)** The precise position of pRNAs transcription initiation in the 6S initiation bubble was mapped based on differential read coverage from (A). The structure of 6S RNA was mapped based on the Rfam annotation RF00013.

**Figure 3.**
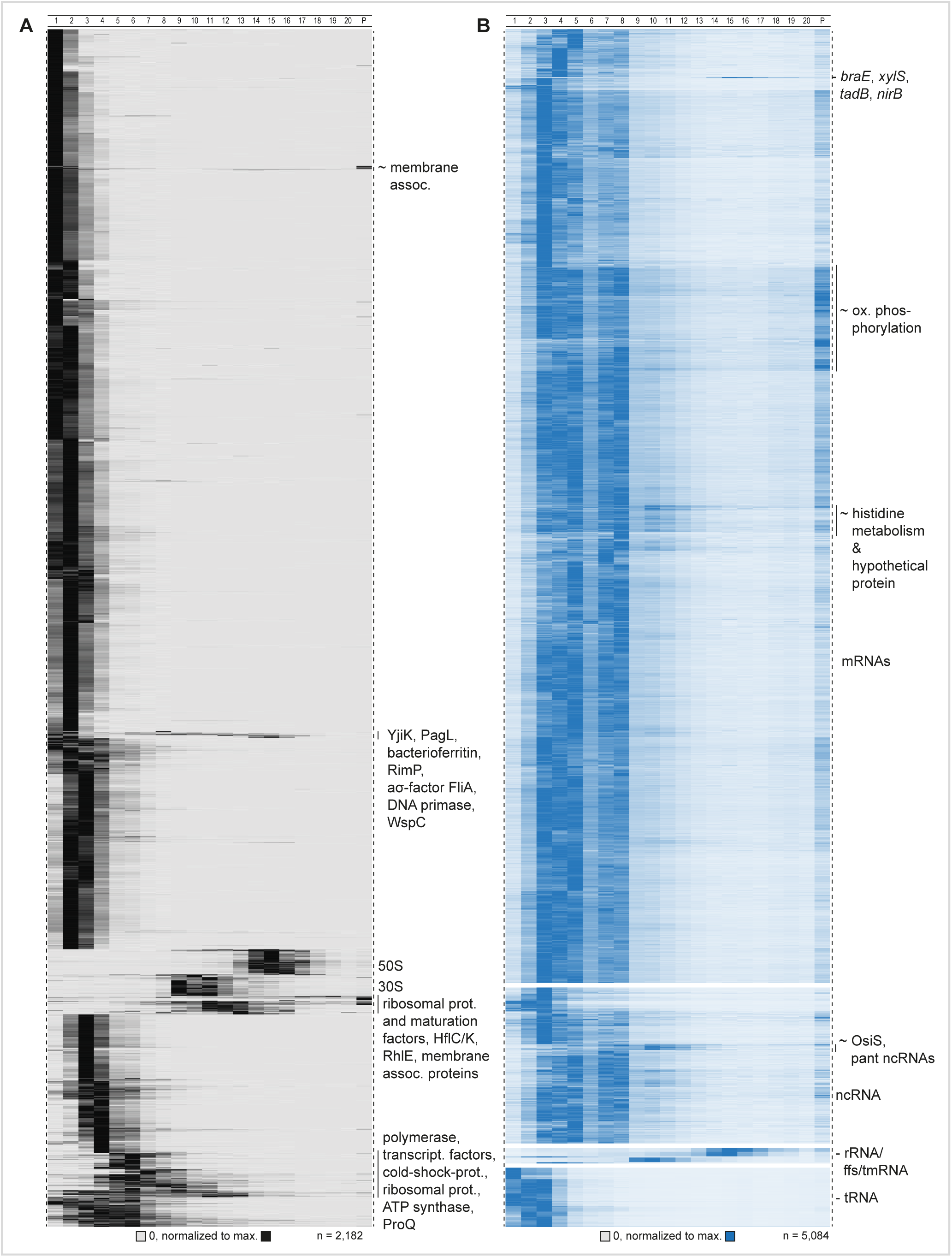
Grad-seq pipeline and all detected sedimentation profiles. **(A)** Proteins sedimented mainly in the first couple of fractions. Defined clusters appeared in fraction below 10. Ribosomal proteins sedimented around fractions 10 and 16. **(B)** Transcripts (merged CDS and UTRs) sedimented mainly in RNAP and ribosomal fractions. tRNAs did not enter the gradient and ribosomal RNAs allocated to HMW fractions.

Viewing the RNA-seq results from individual fractions (**Fig. 3B**), we observed the expected sedimentation of rRNAs in the HMW part of the gradient, whereas tRNAs were found in the LMW top fractions, both of which recapitulated the preceding gel analysis (**Fig. 1D**). However, most of the 5,084 transcripts detected in total sedimented in the first eight fractions, and this equally applied to mRNAs and non-coding RNAs (ncRNAs). With respect to mRNAs, this implies that most of these reads stem from transcripts that are not undergoing translation. However, there is a group of mRNAs with abundant reads in the pellet, indicating association with 70S ribosomes or even polysomes. These transcripts show enrichment for functions in oxidative phosphorylation, the tricarboxylic acid (TCA) cycle, and other metabolic pathways. In essence, these sedimentation profiles provide a glimpse of translational utilization of cellular mRNAs in the exponential growth phase in rich media.

### P. aeruginosa proteins and their complexes

Grad-seq will primarily recover cytosolic proteins as it is performed on a cleared lysate. Indeed, membrane-associated proteins were underrepresented among the ∼2,200 detected proteins (**Fig. 4A**). For a quick overview of functional relationship of co-sedimenting proteins, we clustered all detected proteins by principal component analysis (PCA) according to their sedimentation profiles (**Fig. 4B**). Central regions in the plot correlate with sedimentation somewhere in fractions 4-10, i.e., complexes in the 200 kDa-to 1-MDa range, clear examples of which are the clusters formed by RNAP or ATP synthase subunits. With respect to HMW fractions, ∼6% of the detected proteins show their main peak anywhere between fractions 8 and 20, where they are very unlikely to sediment on their own (**Figs. 3A, 5A**). In other words, most of these proteins are present in stable molecular complexes, indicating that they are associated with a protein or RNA partner to fulfil their function.

**Figure 4.**
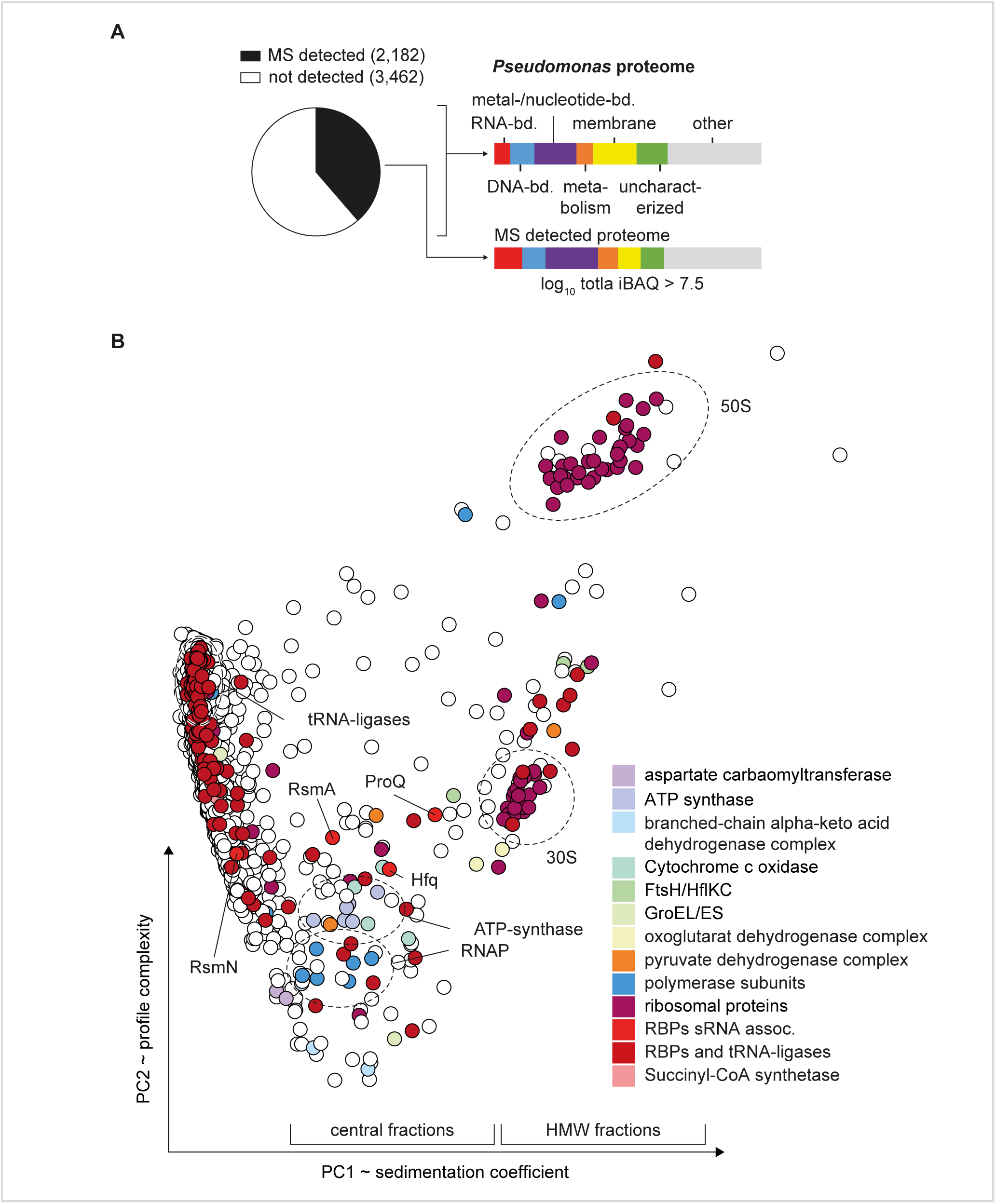
MS detection of complexes and RNPs. **(A)** 2,182 proteins were recovered above the threshold of total iBAQ > 7.5 that represents about 40% of the total proteome. Membrane proteins (yellow) were depleted in the analysed soluble fraction. **(B)** 21 fraction sedimentation profiles were reduced in dimensionality to two principal components (PCs) resembling sedimentation complexity and coefficient. In the central part RNAP and ATP-synthase complexes cluster in proximity with major RBPs Hfq, ProQ, and RsmA/N.

**Figure 5.**
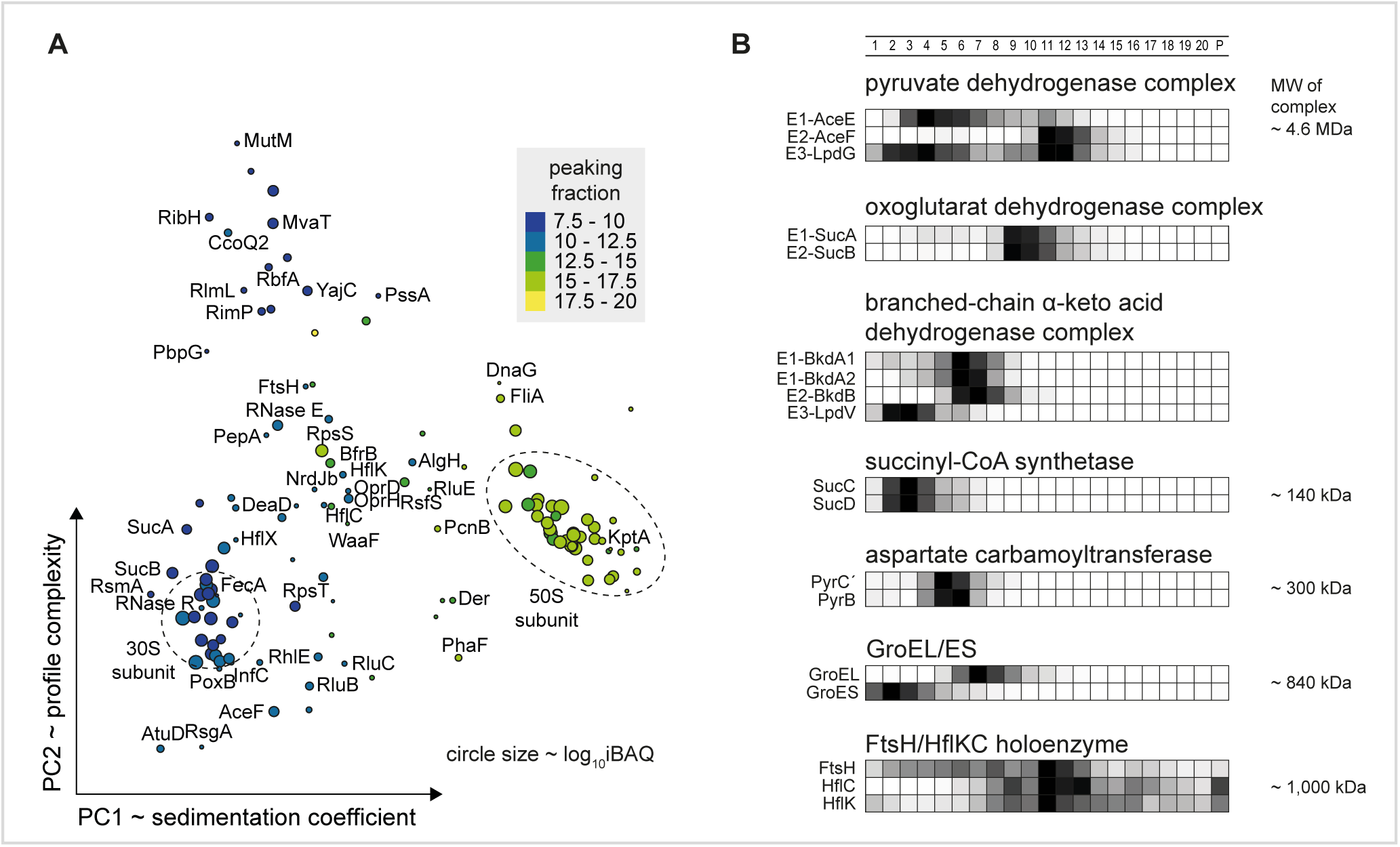
Large complexes co-sediment in higher-molecular weight fractions. **(A)** Sedimentation profiles of proteins that peaked in fractions 8-20 were clustered and labelled by protein name. Dashed circles indicate 30S and 50S ribosomal protein clusters. **(B)** Sedimentation profiles of metabolic complexes are listed with indicated calculated masses. The three dehydrogenase complexes sedimented between fractions 6-12. The TCA cycle complexes succinyl-CoA synthase and aspartate carbamoyltransferase sedimented in fractions 3 and 5, respectively. The chaperone GroEL sedimented in fraction 5 and the associated subunit GroES was dissociated and allocated to fraction 2. The protease complex FtsH/HflKC was detected in fraction 11. The relative abundance is ranging from 0-1 and is colour coded in black for proteins and in blue for RNA.

Next, we evaluated the integrity of complex sedimentation by manual inspection of established proteins, incorporating knowledge from other species such as *E. coli*. Many metabolic processes involve multi-enzyme complexes, some of which are giant. Several such large complexes can readily be discerned from our Grad-seq data (**Fig. 5A**), including all three dehydrogenase complexes: the pyruvate dehydrogenase complex (PDC), the oxoglutarate dehydrogenase complex and the branched-chain α-keto acid dehydrogenase complex (**Fig. 5B**). Interestingly, all of these dehydrogenase multi-enzyme complexes sedimented as much smaller complexes than by their suggested multimeric sizes. This observation is in agreement with a recently determined salt-labile structure of the PDC (38). The succinyl-CoA-synthetase α_2_β_2_ complex, formed by the SucC and SucD proteins (39) from the TCA cycle, sedimented in fraction 3 (MW of ∼140 kDa). The aspartate carbamoyltransferase, which catalyses the first step of the pyrimidine biosynthetic process, builds a complex of ∼310 kDa and sedimented in fraction 5.

Examples of large complexes outside metabolic functions include the tetradecameric 840 kDa GroEL complex, here observed in fraction 7 (**Fig. 5B**). Interestingly, the functionally associated GroES complex sedimented away from it, in LMW fractions, as previously observed in *S. enterica* (22). A particularly large particle seems to be formed by the membrane associated FtsH protease, which is a hexamer that goes on to form a 1 MDa holoenzyme with the membrane complex HflKC (40); we observed it in fraction 11. Importantly, we find dozens of additional proteins to sediment in HMW fractions (**Fig. S2A**). Since many of these are functionally uncharacterized, this region of the gradient bears great potential for discovery of new large protein complexes.

### Topology of the Pseudomonas transcriptome

RNA-seq analysis of the gradient fractions revealed 4,643 transcripts above threshold (>100 reads in all fractions), covering 73.6% of the transcriptome (**Fig. S1E**). Quantitatively speaking, 84% of all reads came from rRNA (**Fig. 6A**). The class of tRNA contributed 6.5% whereas reads from mRNAs were represented by ∼9%. The diverse class of ncRNAs contributed 0.4% of all reads, i.e., markedly fewer than in previous Grad-seq of enteric bacteria in early stationary phase (for example, ∼5% in *E. coli* (28)). Note that in *P. aeruginosa*, too, many ncRNA genes are only upregulated as the cells enter stationary phase (41).

**Figure 6.**
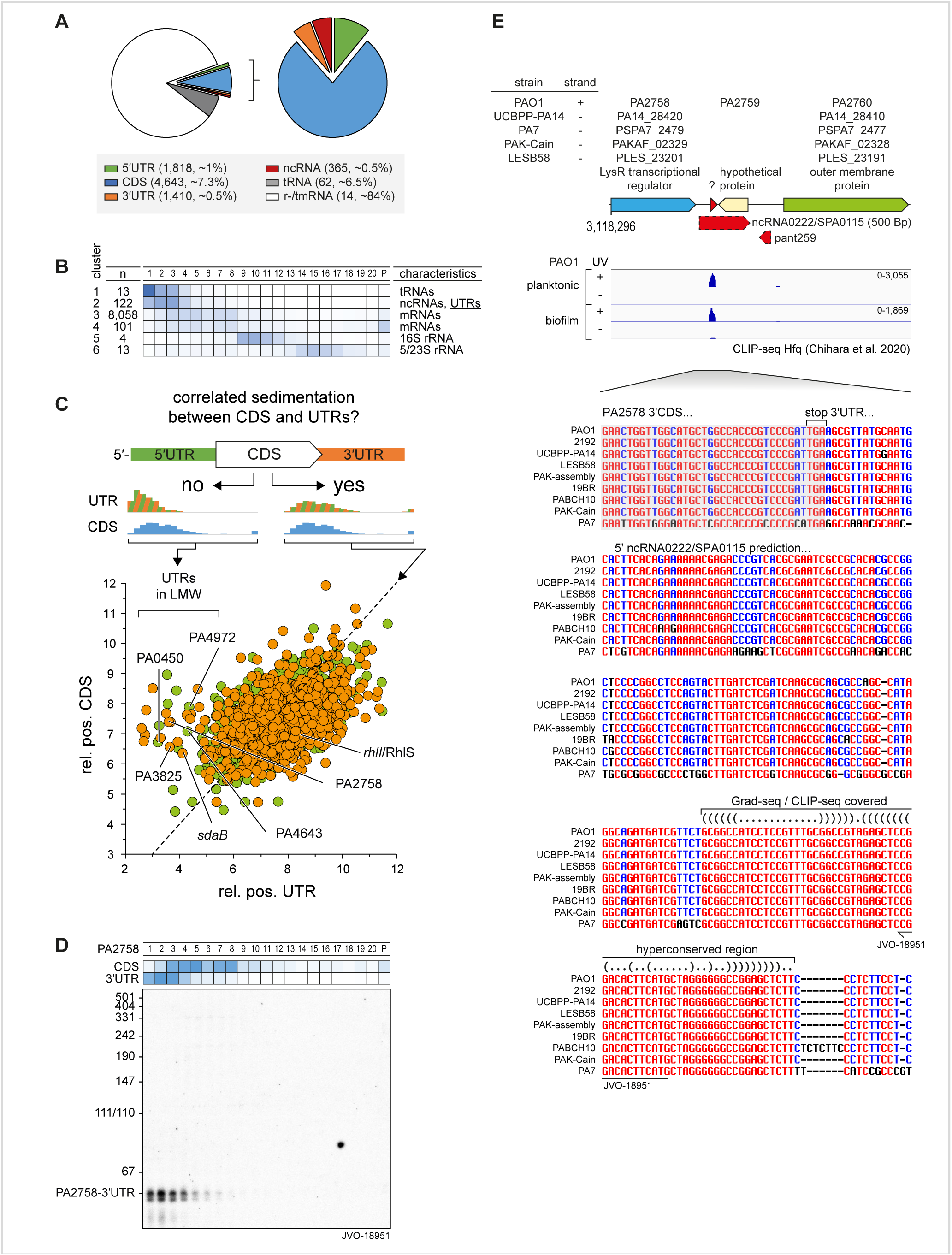
Sedimentation of RNAs. **(A)** 73.6% of *P. aeruginosa* PAO1 transcriptome was detected. mRNAs represented about 9% and ncRNAs about 0.4% sequenced reads. **(B)** RNAs were clustered based on sedimentation position. ncRNAs accumulated together with UTRs in cluster 2. **(C)** Comparison of relative positions of CDS and corresponding UTRs revealed differential sedimentation that attributed to processing and rise of ncRNAs fragments. **(D)** The 3’UTR region of PA2758 was probed by northern for a 50-nt fragment, position was determined previously based on read-coverage in the annotated UTR. **(E)** An alignment of the 3’UTR region of PA2758 revealed a conserved motif in the read-covered part (red arrow). RNA Co-fold (https://e-rna.org/cofold/) resembled a two stemmed structure that is indicated in the dot-bracket annotation.

For a better overview of the general sedimentation of all detected RNA classes, we reduced the complexity of the 21 fractions to six clusters (**Fig. 6B**). The mRNA reads were mainly found in fractions 3-8, coinciding with RNAP, and in the 30S subunit fractions 9-12, as well as in the pellet. This distribution differs from previous Grad-seq analyses in *E. coli*, *S. enterica* and *S. pneumoniae* in which mRNAs generally co-sedimented with ribosomes (22, 28, 29).

Systematic annotation of ncRNAs in *P. aeruginosa* lags behind other model γ-proteobacteria, with only 44 formally annotated ncRNAs (42) as compared to hundreds each in *E. coli*, *Salmonella* and *Vibrio* species (43, 44). However, a total of 473 ncRNAs have been predicted in *P. aeruginosa* (45, 46), 336 of which were detectable in our Grad-seq data. Most of the *P. aeruginosa* ncRNAs and candidates thereof occur in fractions 3-8, thus showing a general sedimentation profile similar to most mRNAs (**Fig. 6B**).

Stable non-coding RNA fragments overlapping with 5’ or 3’ untranslated regions (UTRs) constitute a potentially large class of bacterial riboregulators (47–50). Recent experimental transcriptome annotation (51–53) suggested that also in the *Pseudomonas* strain used here, many such UTR-derived candidate ncRNAs exist and accumulate to considerable levels. Here, we observe that such UTR fragments often show a different sedimentation profile when compared to the respective coding sequence (CDS) of the same mRNA (**Fig. 6C**). These include several UTR fragments that were recently observed to be highly enriched by co-immunoprecipitation with Hfq (8), which is a good indicator of potential cellular function as a base pairing small regulatory RNA (sRNA) (54). Obvious examples are the mRNA 5’UTRs of phosphate transporter PA0450, and the uncharacterized proteins PA4643 and PA4972, or the mRNA 3’UTRs of putative transcriptional regulator PA2758, cyclic-guanylate-specific phosphodiesterase PA3825, and serine dehydratase SdaB, all of which peak at least four fractions away from the CDS part of their parental mRNAs (**Fig. 6C**). With respect to the published literature, RhlS is an activating 5’UTR-derived ncRNA from the *rhlI* gene (55). While the precise mechanism whereby RhlS promotes expression of the *rhlI* mRNA is currently unknown, we notice that the two RNAs occur in different fractions (**Fig. 6C**), supporting the proposed model of an independent regulatory function of the 5’UTR.

To validate the Grad-seq based detection of UTR-derived fragments, we chose to probe the 3’UTR of PA2758 mRNA on a northern blot. We readily detected a distinct ∼50-nt RNA species with the predicted sedimentation in fractions 1-4, distinct from the full-length PA2758 mRNA (**Fig. 6D**). This 3’UTR RNA species was much shorter than the previously predicted ncRNA candidate ncRNA0222/SPA0115 from this genomic region (46). Importantly, however, it comprises the most conserved part of the 3’UTR in question, which includes the predicted Rho-independent transcriptional terminator of PA2758 (**Fig. 6E**). In addition, this ∼50-nt RNA species was recently shown to be highly enriched in global Hfq co-immunoprecipitation experiments ((8); **Fig. 6E**), indicating that it might function as a base pairing sRNA.

*In silico* target prediction using the CopraRNA algorithm (56) supports the notion that the 3’UTR of PA2758 might act *in trans* to regulate mRNAs by base pairing (**Fig. S3**). Focussing on potential binding sites around the start codon of mRNAs, CopraRNA predicts as the top target an extended interaction that might translationally inhibit the mRNA of *napC*, which is the last gene in the *napEFDABC* operon encoding nitrate reductase. Importantly, base pairing would crucially engage a 13-nt long single-stranded region of the PA2758-3’UTR, which looks like a conserved seed sequence of this sRNA. Further studies are required to validate this particular prediction and to determine the targets of the UTR-derived sRNAs proposed here.

### Major RNA-binding proteins and associated ncRNAs

*P. aeruginosa* is known to possess two large post-transcriptional networks that depend on globally acting RBPs: one governed by Hfq, the other by two CsrA-like Rsm proteins. The mRNA and ncRNA ligands of these RBPs have been mapped in a transcriptome-wide fashion (8, 9, 17, 19), albeit not in the growth phase studied here.

The 190-nt PhrS ncRNA, known as a direct activator of the mRNA of the transcriptional regulator PqsR (57), is a well-characterized target of Hfq. PhrS is readily detected in the gradient, by both RNA-seq and northern blot, and shows a very similar sedimentation profile to Hfq (**Fig. 7A**). Its absence from fractions 1-2 suggests that this ncRNA is always present in complex with Hfq and/or other interactors.

**Figure 7.**
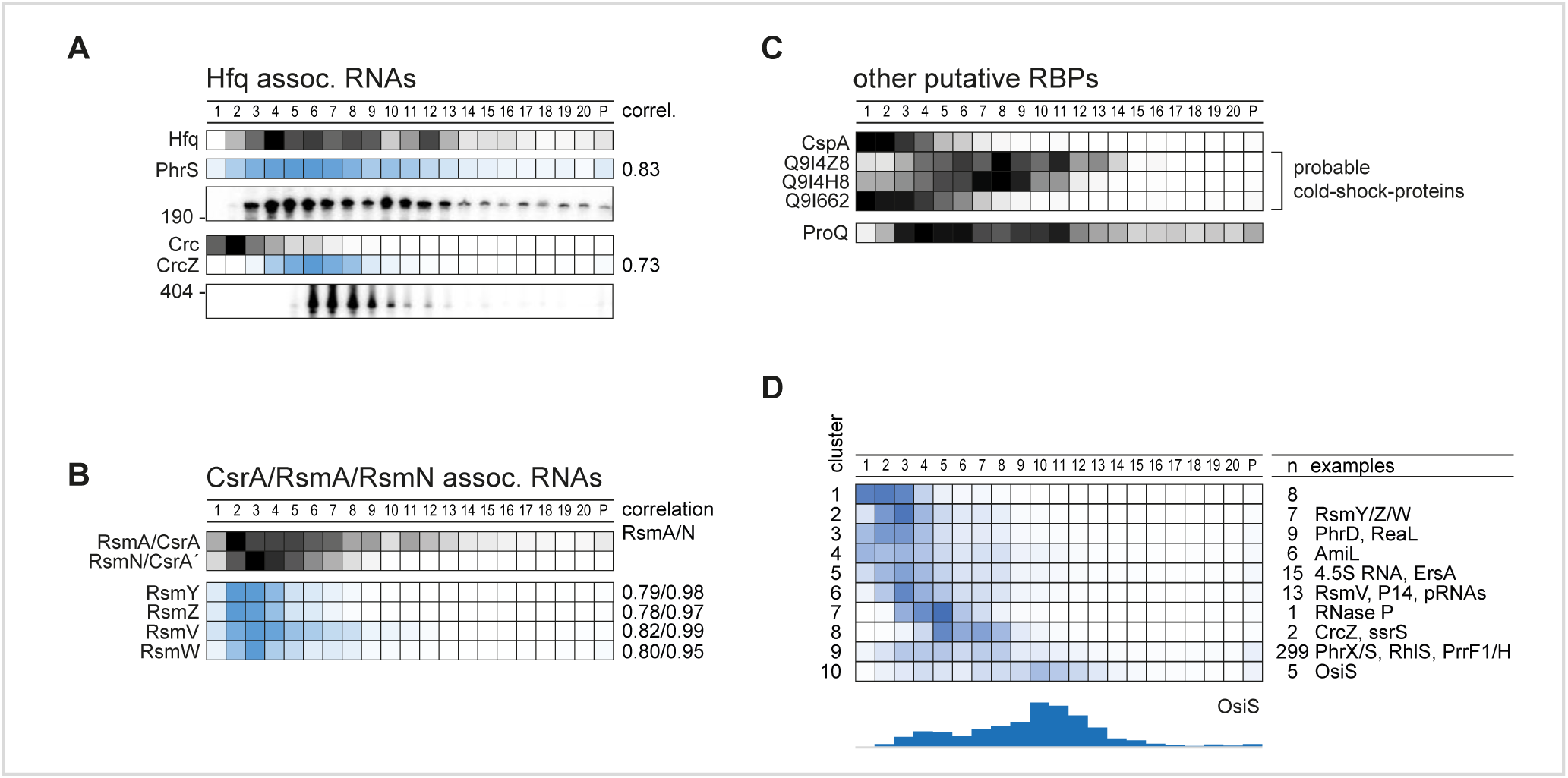
RBPs co-sedimented with targeted RNAs. **(A)** Hfq associated RNAs co-sedimented in a correlated fashion (>0.7) with Hfq mainly in fractions 3-8. **(B)** The same holds true for Rsm RNAs and the corresponding RBPs RsmA and RsmN. **(C)** Other putative RBPs are the cold-shock protein CspA and three putative Csps that sedimented in the first fractions as also previously reported. The FinO-domain protein ProQ sedimented interestingly in between fractions 4 and 11. **(D)** Clustering of annotated ncRNAs by sedimentation position revealed clusters that could be bound by novel RBPs. Interestingly, 5 ncRNAs sedimented in 30S subunit fractions, e.g. OsiS.

Another well-characterized Hfq-related ncRNA is CrcZ. This ∼400-nt ncRNA forms a stable complex with Hfq and Crc (12), which prevents the Hfq/Crc complex from repressing translation of *amiE* mRNA (58). The CrcZ RNA sediments in fractions 6-9 (**Fig. 7A**), in which Hfq is also found. Indeed, the in-gradient profiles of both the PhrS and CrcZ ncRNAs are well-correlated with that of Hfq (coefficient >0.7), suggesting a stable molecular interaction (**Fig. 7A**). By contrast, the Crc protein peaks far away, in LMW fraction 2, suggesting that the Hfq/Crc/RNA complex is unlikely to exist in the rich media conditions used here (as predicted previously; (13)). Additional explanations include that the Hfq/Crc targets are rapidly degraded, leading to disassembly of the Hfq/Crc/mRNA complexes, or that these complexes are simply too labile in a glycerol gradient.

Regarding the Rsm network, the RsmA protein alone is known to regulate —directly or indirectly— 9% of the transcriptome (7, 19, 59). RsmA blocks translation by binding to GGA motifs in mRNAs; its own activity is also regulated at the RNA level, by the ncRNAs RsmY/Z/V/W, which sponge RsmA by presenting multiple GGA motifs (60). The homologous RBP RsmN acts and is acted upon in the same fashion (15, 17, 18). Here, we observe well-correlated sedimentation of the two Rsm proteins and its four-known decoy RNAs, suggesting that in exponentially growing *Pseudomonas* cells (**Fig. 7B**), these two RBPs are largely kept inactive by their ncRNA antagonists. The more abundant RsmA also sedimented in ribosomal fractions, perhaps in association with polycistronic mRNAs. In any case, the RsmA/N profiles are clearly distinct from Hfq and its associated RNAs.

Other RBPs and candidates thereof with unknown ligands also show intriguing sedimentation profiles. Proteomics detected the cold-shock-like protein CspA in the first fractions (**Fig. 7C**), which is in agreement with Grad-seq profiles of CspA in *E. coli* and *Salmonella* (22, 28). Cold-shock proteins, while interacting with many cellular transcripts, bind their targets with low affinity (26), which may explain their accumulation in LMW fractions. Intriguingly, while one of the three other CspA-like proteins of *P. aeruginosa* shows a congruent sedimentation profile, the other two migrate towards the middle of the gradient. Thus, these two putative RBPs, PA0961 and PA1159, represent strong candidates for cold-shock proteins that engage in stable complexes with either RNA or other proteins (**Fig. 7C**).

Chromosomally encoded RBPs with a FinO domain such as ProQ have recently emerged as the third major ncRNA-associated RBP in enteric bacteria (25). Based on a Pfam prediction (61), we propose protein Q9I0Q4 as the putative ProQ homolog in *P. aeruginosa*. Peptides of it can be detected in almost all gradient fractions, although the majority of them are found in fractions 3-11 (**Fig. 7C**). This ProQ profile is broader than in *E. coli* or *Salmonella*, especially with respect to *P. aeruginosa* ProQ appearing in 30S ribosome fractions. In any case, this broad sedimentation argues for *P. aeruginosa* to possess functional FinO/ProQ-like RBPs.

To obtain a focused overview of ncRNA sedimentation patterns for potential cross-comparison with RBPs, we clustered all 365 detected and candidate ncRNAs according to their sedimentation profile, yielding ten clusters (**Fig. 7D**). Approximately three-quarter of these ncRNAs end up in cluster 9, which includes the well-characterized Hfq-associated ncRNA PhrS and PrrF1 (57, 62), and PhrX (63). In essence, many of these ncRNAs sediment like mRNAs (**Fig. 6B**, cluster 4). This is particularly pronounced with the five members of cluster 10, challenging the previous annotation of these transcripts as noncoding RNAs. For example, OsiS (PA0611.1, pant67, (45)) has been suggested to act as a regulatory RNA to link oxygen level sensing to the production of quorum sensing molecules (64). However, the co-migration of these transcripts with 30S ribosomes suggests the possibility that they possess overlooked small ORFs. Note that a putative 30S association recently guided the discovery of a functional small ORF in the seemingly noncoding RyeG RNA of *E. coli* (28, 65). In the present case, recent *P. aeruginosa* Ribo-seq data (66) provide additional support for our prediction that OsiS is translated into a protein (**Fig. S4**).

### Phage-induced stress affects cellular organization of the RNome

The in-gradient distributions of RNA and protein described above reflect the complexome of *P. aeruginosa* under optimal growth conditions, characterized by rich supply of nutrients and rapid growth. To determine how this molecular organization is perturbed by a biotic stress, we subjected *P. aeruginosa* to infection by a phage. Specifically, the lytic giant phage ΦKZ which is a well-established model phage of pseudomonads and known to usurp many cellular functions of its host (67–69) (**Fig. 8A**).

**Figure 8.**
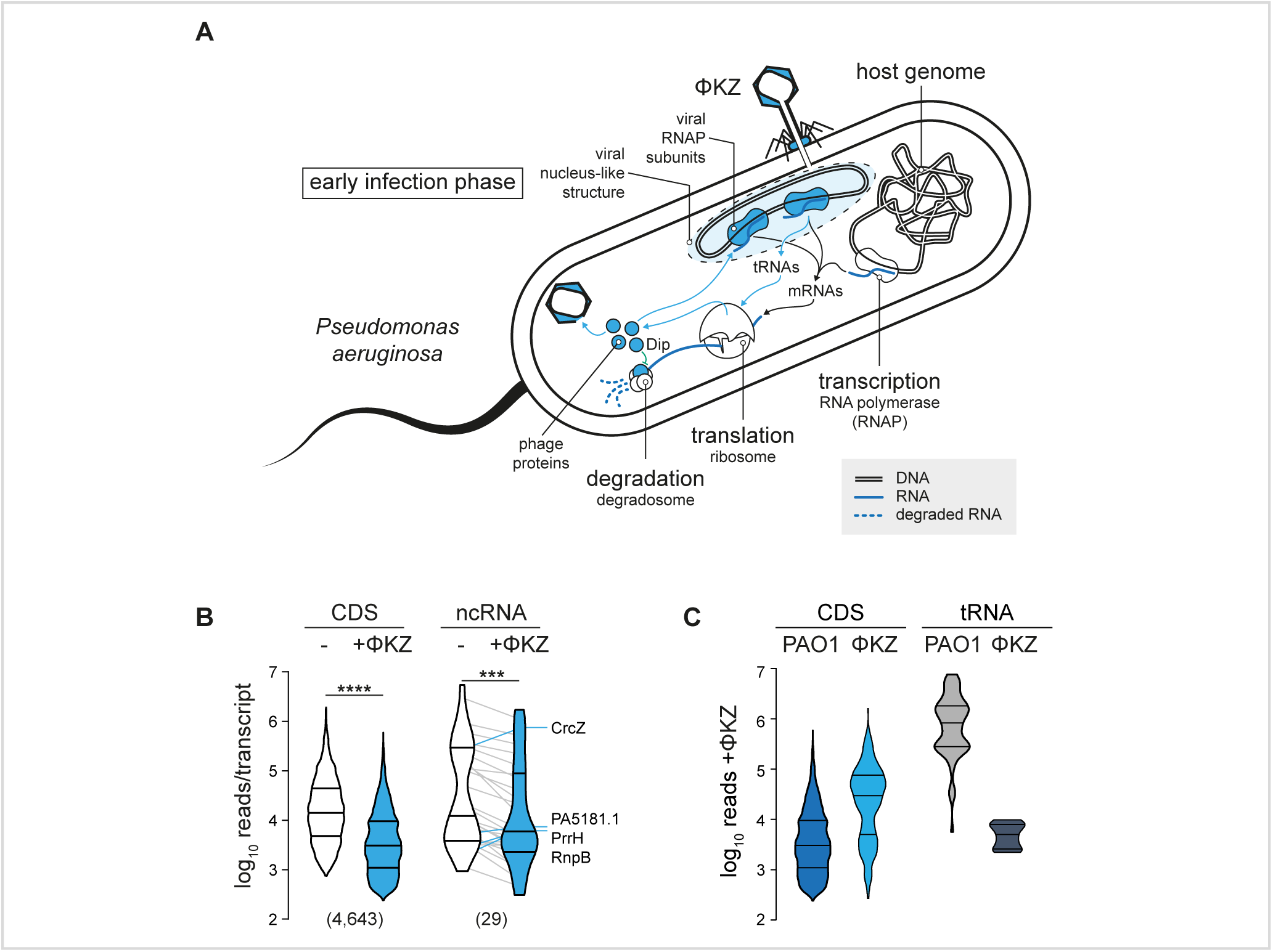
Host RNome is altered at early infection. **(A)** ΦKZ conquering of *Pseudomonas* is strongly linked to modulation of RNA homeostasis. Viral RNAPs are utilized and produced to transcribe phase genes. Additional phage tRNAs are utilized in translation, and the RNA degradation is blocked through the Dip protein. **(B)** Host CDS and ncRNA levels were significantly reduced by about 5-fold in a non-parametric paired t-test. The threshold for statistical significance was a two-tailed P value of <0.05 (\*\*\**p* < 0.001; \*\*\*\**p* < 0.0001). Prism 9 was used for statistical analyses. Lines connect individual RNA levels. Outliers that increased levels upon ΦKZ infection are labelled. **(C)** Individual phage CDS are two magnitudes more abundant than host RNAs, whereas phage tRNAs remain underrepresented 10 min post infection.

Performing Grad-seq on *P. aeruginosa* cells after 10 min of infection by phage ΦKZ, we found that phage transcripts accounted for ∼3% of all reads, while the host mRNAs remained at ∼7% (**Fig. S5**). Nonetheless, a paired t-test showed that the sum of all host CDS and ncRNAs at this early infection time point was approximately fivefold lower compared to uninfected cells, notwithstanding a few prominent outliers such as the ncRNAs CrcZ, PrrH, and RnpB (**Fig. 8B**). At the same time, the individual ΦKZ transcript exceeded the typical host transcript in abundance by one order of magnitude (**Fig. 8C**).

While altered abundance of host transcripts is one consequence of phage predation, we were particularly interested in the possibility that the phage infection might also perturb host transcriptome organization, altering the association of host transcripts with cellular factors to promote viral replication. Such changes, which should be visible as shifts in the gradient, could point out stress response mechanisms and pathways that remain invisible in standard RNA-seq experiments.

Focussing on transcripts with a <3-fold change in total level, we indeed observed hundreds of transcripts with negative or positive shifts in their sedimentation profile (**Fig. 9A,B**). An excellent example is given by the mRNAs of the uncharacterized proteins PA4441 and PA2746a, which both showed redistribution towards fractions 10-11, possibly reflecting increased translational initiation by 30S ribosomes. Another example is *liuR*, coding for the transcriptional regulator of the of metabolic *liu* genes (70) (**Fig. 9A,C**). Similarly, we predict several host ncRNAs to reassociate with different partners in the early phase of phage infection, illustrative examples of which are ErsA, pant512 and pant253 (**Fig. 9D**). By contrast, with the exception of tRNA^Leu^, transfer RNAs generally stayed in the same fraction after ΦKZ infection (**Fig. 9A-C**).

**Figure 9.**
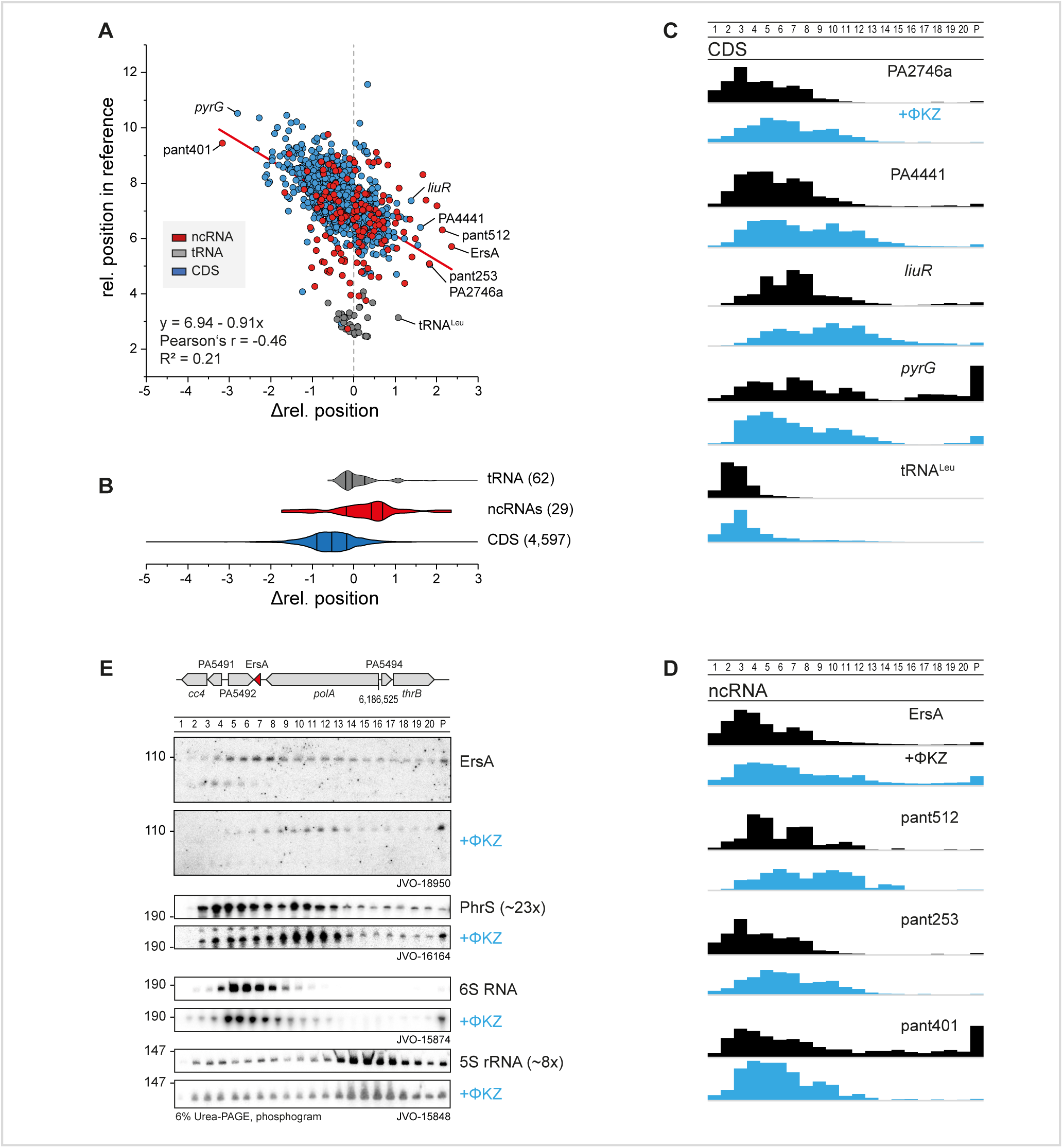
ΦKZ infection shifts transcripts and ncRNAs. **(A)** Transcripts that sediment in HMW fractions are more shifted towards LMW fractions. (change in level <3-fold). **(B)** Annotated ncRNAs were shifted in direction of HMW fractions. **(C)** Sedimentation profiles of host CDS that shift strongly towards H/LMW fractions from (A). **(D)** Sedimentation profiles of phage-infection affected host ncRNAs, of which ErsA represents an important regulator in biofilm formation and is shifted to HMW fractions. **(E)** The ErsA shift in sedimentation was validated by northern probing against the end of stem-loop I in ErsA and accumulated strongly in the pellet fraction. A less pronounced shift was observed for PhrS that was strongly depleted upon infection.

ErsA is a well-characterized small RNA of *P. aeruginosa*, which has been proposed to serve a dual regulatory role in biofilm development, activating the mRNA of transcriptional regulator AmrZ while at the same time repressing the mRNA of AlgC, an important virulence-associated enzyme for the production of sugar precursors (71, 72). In addition, ErsA acts as a translational repressor of the mRNA of outer membrane porin OprD (73). ErsA is well-expressed during the exponential growth phase (51) and bound by Hfq (8), making it an excellent candidate for validation by an independent technique of the infection-dependent changes seen in Grad-seq. Using an oligonucleotide probe for ErsA, we probed all 20 gradient fractions plus the pellet by northern blotting. Intriguingly, this method showed even more clearly that EsrA, 10 min into phage infection, is depleted in LMW fractions, with its main peak shifting to fractions 10-11 where 30S ribosomes sediment (**Fig. 9E**). A similar, albeit less pronounced shift was observed for the PhrS ncRNA. By contrast, phage infection did not affect the in-gradient distribution of 5S rRNA and 6S RNA, as predicted by the digital data. While the biological meaning of the redistribution of EsrA remains to be understood, its independent validation by northern blot adds confidence for the predicted infection-induced shifts of other cellular transcripts.

### Phage mRNAs are efficiently transferred from transcription to translation

At the early time point of ΦKZ infection used here, nearly all (366/376) annotated phage genes (NC_004629.1, NCBI nucleotide) are expressed (**Table S1**, (30, 51)). For an initial glimpse at the sedimentation profiles of phage transcripts, we probed two well-expressed viral operons on northern blots; specifically, operons 54 and 88 (30), which encode metabolic and structural proteins of the phage, respectively. Intriguingly, in both cases the strongest signal obtained were from the pellet fraction, indicating that these mRNAs are heavily engaged by the translation apparatus (**Figs. 10 and S5F,G**). In addition, signals within the gradient were primarily seen in fractions 10-13, overlapping with the 30S ribosome peak.

**Figure 10.**
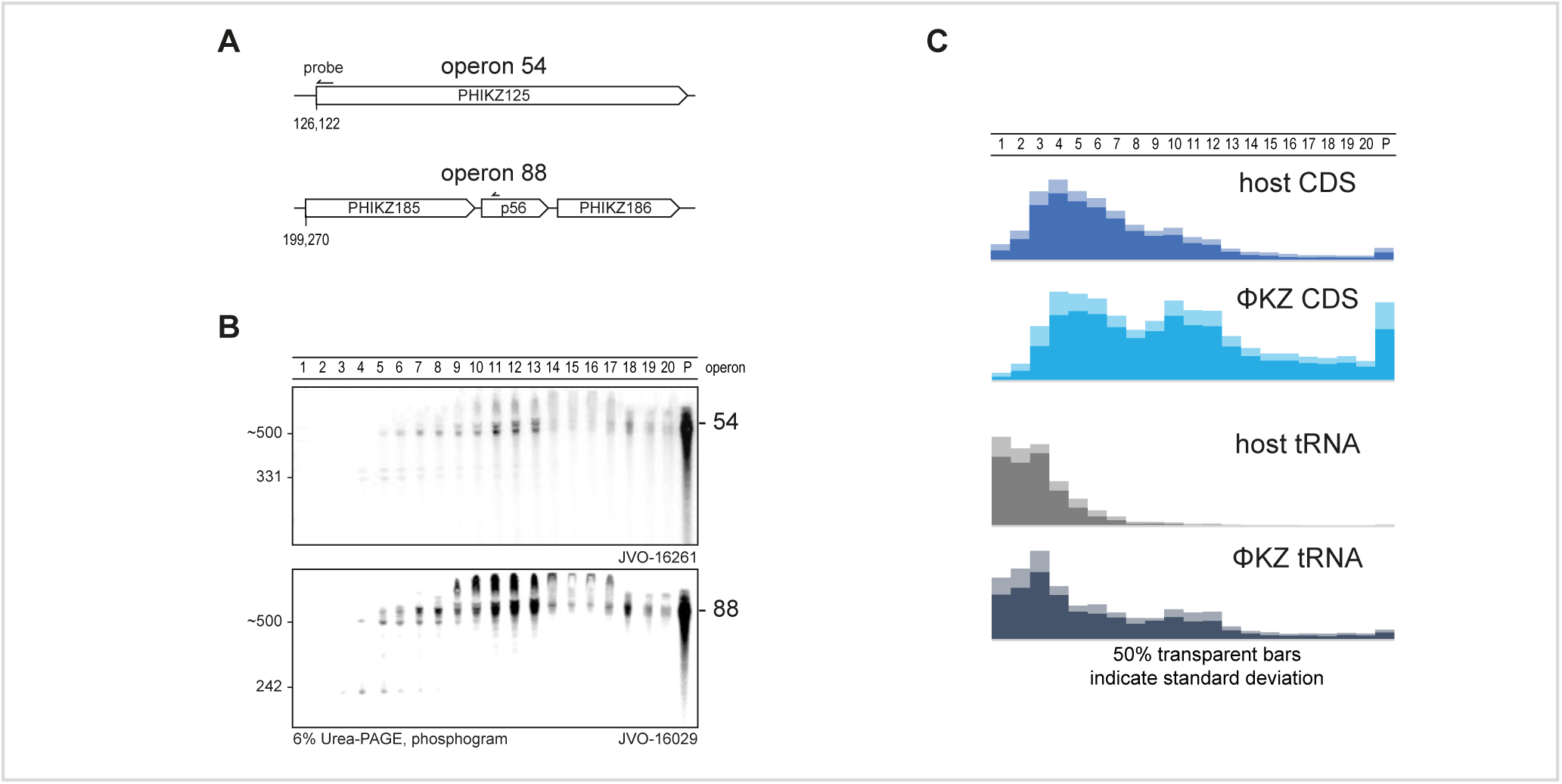
Phage RNAs associate strongly with ribosomes. **(A,B)** Expression and sedimentation of early infection operons 54 and 88 was northern probed and revealed a strongly ribosome associated profile. Interestingly, smaller fragments sedimented exclusively before fraction six. **(C)** In general, phage transcripts sedimented substantially at ribosomal fractions indicating an efficient transfer to translation or stabilization at the ribosome. Phage tRNAs were also strongly utilized in translation or stabilized at ribosomal subunits judged by sedimentation in ribosomal fractions.

Global inspection of the Grad-seq data reinforced the notion that phage mRNAs are prioritized for translation. That is, in contrast with host mRNAs which were primarily detected in fractions 3-8 (**Fig. 6B**), the phage mRNAs were clearly more abundant in fractions containing 30S and 70S ribosomes (**Fig. 10B,C**). In addition, different from host tRNAs, the phage-encoded tRNAs also appeared in the 30S-containing fractions (**Figs. 6B, 10C, S5E,F**), which appears to be a type of association (74). Thus, two classes of ΦKZ transcripts indicate the existence of mechanisms that ensure efficient synthesis of phage proteins.

### Grad-seq guides the identification of phage-encoded ncRNAs

We reasoned that if phage mRNAs generally co-migrate with ribosomes, non-coding phage transcripts might be revealed through different in-gradient distributions, as already evident from the phage-encoded tRNAs. Indeed, manual inspection of Grad-seq profiles along the 280-kbp phage genome revealed two ncRNA candidates. The first example was a ∼100-nt RNA species from the intergenic space between phage genes PHIKZ298 and PHIKZ297, predicted to sediment primarily in fractions 3-4 (**Fig. 11A,B**). Northern blot probing proved this species to be abundant and also confirmed its predicted length. Interestingly, the probe used here simultaneously detects a larger transcript(s) with an mRNA-like in-gradient distribution. Combined with the lack of a mapped TSS downstream of the stop codon of PHIKZ298 (51), we assume that the detected 100-nt RNA species is generated by 3’UTR processing of the PHIKZ298 mRNA. According to *in silico* prediction, the ncRNA forms two stem-loops, the second of which might constitute the Rho-independent transcriptional terminator of the PHIKZ298 gene (**Fig. 11C**).

**Figure 11.**
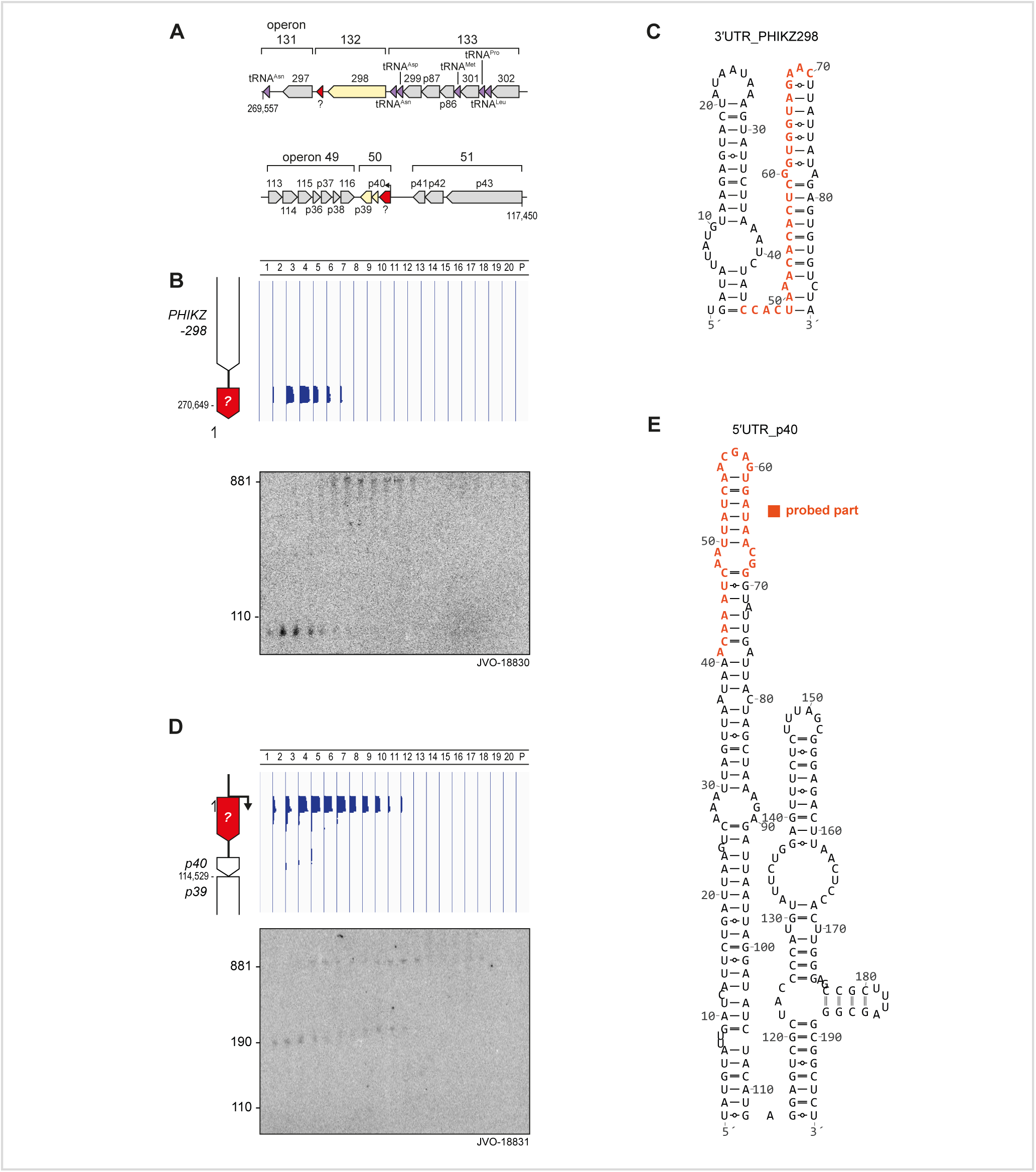
Grad-seq guides the discovery of phage-related ncRNA candidates. **(A)** Loci of operons 132 and 50 including candidate ncRNAs. **(B)** In the 3’UTR of PHIKZ298, we observed sedimentation exclusively around fraction four that was confirmed by probing a ∼100-nt fragment that M-folded (108) into a two stemmed structure **(C)**. **(D)** Read-coverage profiles in the 5’UTR of p40 revealed an LMW weighted sedimentation profile different from phage CDS transcripts (Fig. 10C). A ∼180-nt fragment was detected in RNAP associated fractions and at 30S subunit fractions. A longer fragment (∼880-nt) was substantially shifted to fraction 12. **(E)** The 5’UTR-p40 fragment M-folded into a two stemmed structure. Northern probing was conducted against the nucleotides indicated in orange.

The other ncRNA candidate also accumulates in the LMW part of the gradient, yet shows a broader distribution (fractions 4-10; **Fig. 11A,D**). This putative ncRNA comprises the first 180-nts of the 5’UTR of phage gene p40. Similar to the above candidate, northern blot probing confirmed both the proposed length of the candidate ncRNA itself and detected larger transcripts with mRNA-like behaviour in heavier fractions. Note that for reasons unknown, the signals of both these putative ncRNAs are shifted by one fraction towards the top of the gradient when comparing the northern blot to the Grad-seq data. The 5’UTR-derived ncRNA species also seems highly structured, showing two stem loops (**Fig. 11E**), the second of which might promote premature termination before transcription reaches the start codon of p40.

We carefully checked both UTR-derived RNA species for potential small ORFs, including alternative start codons, but found none that would be preceded by a strong Shine-Dalgarno sequence. We therefore consider both as ncRNAs and take them as illustrative examples how the extra information provided by Grad-seq, as compared to simple RNA-seq, may help to discover overlooked transcripts in phage genomes.

## DISCUSSION

The primary objectives of this work have been: i) to provide the interested microbiologist with previously unavailable Grad-seq profiles of cellular transcripts and proteins in the model bacterium *P. aeruginosa* from a commonly-studied growth condition, ii) to assess the quality of these high-throughput results by independent methods, iii) to illustrate how the technique might be utilized in the future to study bacteriophage-host interactions, and iv) to organize the data such that they can be accessed and interrogated by non-specialists. To facilitate the latter, we have set up an online browser (**Fig. 12**), which is available https://helmholtz-hiri.de/en/datasets/gradseqpao1/ and allows the user to cross-compare the present *Pseudomonas* data with Grad-seq results for several other species (22, 28, 29). In addition, the browser offers comparison with GradR (75), which is a recently developed variation of Grad-seq trying to predict potential RNA association/dependency of cellular proteins, and more specifically, unrecognized RBPs. Although those GradR data were obtained with *Salmonella*, some of the many proteins affected by the RNase treatment used in this method will also be conserved in *P. aeruginosa*.

**Figure 12.**
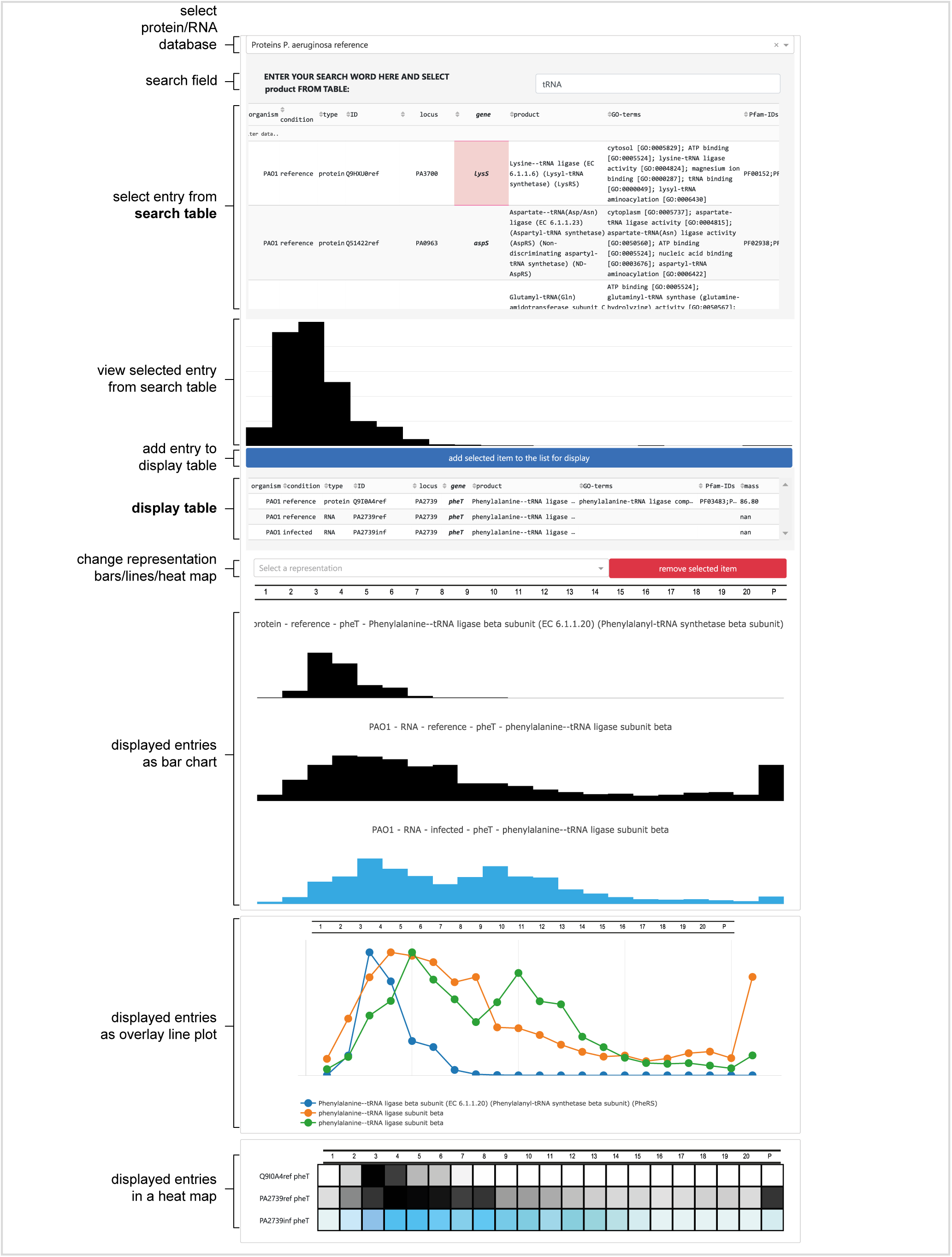
Differential Grad-seq browser. Protein and RNA sedimentation profiles can be searched for non-infected and infected experiments. Selected entries are displayed as bar-chart and can be added in a display list that is plotted as bar-chart, lines, or heatmap. The browser is accessible at https://helmholtz-hiri.de/en/datasets/gradseqpao1/

Even though the present *P. aeruginosa* Grad-seq data covers only one growth condition, we believe that it already offers a huge discovery space. Given the bacterium’s impressive metabolic capabilities, the MS-profiles of all detected proteins should give new insights into metabolic multi-enzyme complexes. Obviously, not all these complexes are sufficiently stable to survive the long (over-night) ultra-centrifugation procedure during which cellular complexes separate according to size and shape in a glycerol gradient. For example, the PDH complex dissociated into stable smaller subcomplexes (**Fig. 5B**), which is in keeping with recent size-exclusion experiments that revealed a salt-labile structure of the macromolecular complex (38). Nonetheless, in addition to the examples highlighted in **Fig. 5B**, there are many proteins that sedimented between the fractions of the small and large ribosomal subunits. In other words, these proteins must be part of stable complexes in the 2-4 MDa range, an information that should provide useful leads for future functional analysis.

From an RNA biology perspective, the initial Grad-seq data provided here promises to be a treasure trove. Known RBPs such as Hfq and RsmA/N sedimented in a much higher molecular weight range than their individual size would predict. We have recently shown for Hfq (and other RBPs) in *Salmonella* that this sedimentation behaviour is primarily due to RNA interactions. Indeed, when transcripts are removed by RNase treatment of the lysate, these proteins shift to LMW fractions (75). In the present *P. aeruginosa* data, the associations of the Hfq and RsmA/N RBPs are also obvious by a clear co-migration with their major RNA ligands (**Fig. 7A,B**). While these three RBPs have seen extensive work, our Grad-seq data strongly indicate that there is yet another global RBP. That is, we have detected peptides of a candidate ProQ RBP (protein PA2582), which is encoded in a chromosomal region close to exonuclease repair gene *uvrC* and the regulatory protein PA2583 (**Fig. S6**). The predicted ProQ of *P. aeruginosa* not only carries a full-length FinO domain, it also shows conservation of several important residues for RNA binding by *E. coli* ProQ (76). The pseudomonal ProQ proteins are small (∼20 kDa), and similarly to *Neisseria* ProQ (21), lack the typical C-terminal or N-terminal extensions of the central FinO domain (25), which makes them attractive candidates for understanding the core recognition mode of RNA by the FinO domain.

The complex of 6S RNA and RNAP is an RNA-based regulation previously undescribed in *P. aeruginosa*. The two partners show in an almost ideal form both, narrow peaks and exceptionally high correlation in the gradient, primarily in fractions 5-6 (**Fig. 2A**). While similarly high correlations have been shown before in both conventional and Grad-seq based analyses of 6S RNA-RNAP (22, 28, 29), the new observation in the present *P. aeruginosa* Grad-seq experiment are the 14-nt pRNAs. These pRNAs are a natural product of the 6S RNA regulatory cycle in which 6S RNA captures RNAP by mimicking an open promoter complex of transcribed DNA, and once bound, changes the affinity of RNAP for different σ factors, the result of which is a global change in transcription (34). In order to dissociate from 6S RNA, RNAP uses it as a template for transcription, yielding pRNAs. Thus far, these pRNAs have been considered as an inevitable side product of 6S RNA release, possessing no independent function and awaiting degradation. However, our detection of a pRNA peak spanning several fractions (2-5) may now hint that these short transcripts go on to interact with other cellular molecules, perhaps as part of serve a currently unrecognized independent function in the cell.

Next to Grad-seq analysis of naïve *P. aeruginosa* cells, we provide an RNA snapshot with the corresponding experiment performed at an early time point of phage infection. Phages by definition manipulate host cell metabolism in exquisitely diverse ways, as reviewed recently (68). Specifically, *Pseudomonas* bacteriophage ΦKZ is known to rapidly shift towards host-independent viral replication. This is achieved through transcription by both, virion-delivered and virus-encoded RNAPs (30), a complete reorganization of the metabolism (77), and modulation of the host RNA turnover machinery (78). However, insights into RNA/protein interactions between phage and host remain scarce. Therefore, the RNA data provided in the next section allow a first glimpse at how ΦKZ alters RNA processing in the host, and hint at modulation of translation rates of some host mRNAs, and of phage mRNAs in general. Importantly, active manipulation of protein synthesis has been reported for coliphages T4 (79) and T7 (80–84), and is also implied by prediction of phage-encoded S1 proteins (85, 86).

On the host side, phage infection impacts the in-gradient distributions of both mRNAs and ncRNAs. Many host mRNAs shift away from ribosomal fractions, and this effect is even more pronounced for those transcripts with a particular strong ribosome association in the first place. Primary examples include transcripts related to oxidative phosphorylation (**Fig. S7**); a strong reduction of the synthesis of these proteins may eventually lead to a collapse of cellular energy production. In general, given that active translation stabilizes bacterial mRNAs, the predicted interruption of translation may offer one explanation for the observed ∼5-fold depletion of host transcripts 10 min after phage infection.

Yet, we also observe the reciprocal behaviour, i.e., host mRNAs that move into ribosome-associated fractions. A particularly noteworthy case is the mRNA of protein PA4441, which carries a predicted DUF1043 domain that is also found in cyclic-oligonucleotide-based anti-phage signalling system (CBASS) effectors (87). Put simply, after phage predation, the mRNA of a potential phage defence protein migrates to the 30S-related fraction as the consequence of increased translational initiation. This may indicate the existence of a specific mechanism to rapidly activate synthesis of host proteins with protective functions.

In addition to mRNAs moving out of or into ribosomal fractions, Grad-seq predicts interesting cases of ncRNAs that seem to associate with different or additional cellular partners after lytic infection. These shifts for the ErsA and PhrS sRNAs were independently validated on northern blots (**Fig. 9E**). Because both these ncRNAs are well-characterized, they would serve as ideal models to understand the mechanisms behind the observed shift. In the case of PhrS, it is worth mentioning that in *P. aeruginosa* strain PA14 this sRNA may promote adaptive immunity against bacteriophages through suppression of Rho-dependent termination of early CRISPR transcripts (88). This CRISPR leader is conserved in the PAO1 strain used here despite the lack of a CRISPR array, and we thus speculate that the observed shift of PhrS RNA might be the result of enhanced PhrS-CRISPR leader interaction upon phage infection.

Of well-understood molecular mechanisms whereby phages usurp bacteria, several include active manipulation of host RNA decay (89, 90). The present Grad-seq data is of insufficient depth to allow for a statistically sound global analysis of phage-induced alteration of host RNA processing. However, during inspection of rRNA profiles, we observed that biogenesis of 5S rRNA in phage-infected cells might be compromised, as suggested by a smeary northern blot signal for 5S rRNA in several gradient fractions (**Fig. S8A**). Processing of 5S rRNA from longer ribosomal transcripts crucially depends on the major endoribonuclease RNase E, and ΦKZ expresses a protein called Dip that functionally interferes with the RNase E-associated degradosome (78, 89, 91).

In further support of altered RNase E activity, we observe a differential accumulation of 5’ mRNA fragments of RNase E itself (**Fig. S8B,C**), which is well-established to cleave in the 5’UTR of its own mRNA to feedback-control its own protein levels (92–94). Upon ΦKZ infection, a longer band that we attribute to cleavage at the *rne-II* motif (Rfam RF01756, (95)) disappears. Whether the altered processing eventually causes a decrease in RNase E protein levels needs to be shown. Interestingly, this mechanism would be the opposite to what was reported with the marine cyanobacterium *Prochlorococcus* MED4 in which phage infection boosts RNase E synthesis by altered processing of its mRNA (96).

Finally, one of the most intriguing phage-related observation in our Grad-seq data is the distribution of phage versus host mRNAs. Contrary to mRNAs for which reads where detected in both LMW and ribosome-containing HMW fractions (**Figs. 3B, S5E**), phage mRNAs sediment almost exclusively in the HMW part of the gradient as well as in the pellet (**Figs. 10C, S5F,G**), the latter of which contains 70S monosomes and polysomes. On the face of it, this would suggest that ΦKZ mRNAs are prioritized for translation, perhaps even at the expense of making host proteins, as has been reported previously in other systems (79–86) We would like to caution, however, that mRNA reads in LMW fractions represent a steady-state picture in which we see host mRNAs undergoing degradation, whereas the newly produced phage mRNAs, only 10 min after the phage has attacked, may still be fully protected by ongoing translation. In addition, phage ΦKZ segregates its DNA from immunity nucleases of *P. aeruginosa* by constructing a proteinaceous nucleus-like compartment (69), which might provide further protection of phage mRNAs. While this new model does not question generally that phage mRNAs are translated by cytoplasmic ribosomes, the influence of this intracellular spatial segregation on phage mRNA stability is not clear. Future work with higher time resolution will be needed to determine whether phage mRNAs are indeed prioritized for translation, and if so, which phage or host factors are responsible.

Many of the above observations will benefit from a further mechanistic analysis of the Grad-seq protein data of infected host cells, which is currently ongoing. In studying the molecular mechanisms governing phage-bacteria interactions, it should be obvious that Grad-seq offers unique opportunities to reveal RNA-protein interactions, including phage-CRISPR-Cas complex interactions, usage and modifications of tRNAs or the discovery and functional elucidation of phage ncRNAs, all of which have remained largely unstudied to date. Indeed, the potential of differential sedimentation profiles in a single sample became apparent as we were able to point to two previously unannotated ncRNAs in the phage (**Fig. 11**), whereas TSS mapping in ΦKZ (51) failed to predict such candidates. To put this into perspective, a mere 21 ncRNAs have been described in phages to date (97), hence Grad-seq offers an orthogonal approach for identification of RNA processing sites and ncRNAs.

## MATERIAL AND METHODS

### Cell preparation, lysis, and gradient sedimentation

*Pseudomonas aeruginosa* strain PAO1 (JVS-11761, **Table S2**, German Collection of Microorganisms and Cell cultures GmbH: DSM 22644, BacDive ID: 12801, NCBI tax-ID: 208964) was grown in 400 mL LB media to OD600 0.3 at 37°C. Optionally, the cells were infected with phage ΦKZ at a multiplicity of infection (MOI) of 15 to ensure immediate infection of the entire culture. To ensure that infected cell culture was in the early infection phase, the infected culture was mixed and incubated for 5 min at room-temperature without shaking, followed by incubation at 37°C for 5 min with shaking, as described previously (30). The culture was cooled rapidly down in ice and the cells were pelleted at 6,000 × g for 15 min at 4°C, and washed three times with ice-cold TBS (20 mM Tris/HCl pH 7.4, 150 mM NaCl).

The gradient analysis was performed as published (22), with minor modifications. Cells were resuspended in 0.5 ml lysis buffer (20 mM Tris/HCl pH 7.5, 150 mM KCl, 1 mM MgCl_2_, 1 mM DTT, 1 mM PMSF, 0.2% (v/v) Triton X-100, 20 U/ml DNase I (Thermo Fisher Scientific), 200 U/ml SUPERase-IN (Life Technologies)) and one volume of 0.1 mm glass beads. Lysis was obtained by vortexing this mix for 30 s, followed by 15 s cooling on ice. This process was repeated ten times and the lysate was cleared at 12,000 × g for 10 min at 4°C. The supernatant was used for SDS-PAGE analysis and RNA preparation by TRIzol™ (Thermo Fischer Scientific). Aliquots of 180 μl supernatant were layered on a linear 10-40% (w/v) glycerol gradient and were sedimented in a SW40Ti rotor at 100,000 × g for 17 h at 4°C. The gradient was fractionated in 590 μl fractions and the absorption was determined at 260 nm. Protein samples for SDS-PAGE were prepared for each individual fraction and analysed as previously described (22). A 0.5 ml volume of each fraction was used for RNA isolation by addition of 1% SDS, and subsequently 600 μl acidic PCI (Carl Roth). Samples were vortexed extensively and the phases separated by centrifugation. The upper aqueous layer was extracted repeatedly with chloroform. The aqueous layer was selected and precipitated by three volumes ice-cold ethanol with 0.3 M NaOAc (pH 6.5). GlycoBlue^®^ (1 μl) (Thermo Fischer Scientific) was added as tracer. After overnight storage at -20°C the precipitated RNA was pelleted and washed with 70% ethanol, dried and dissolved in nuclease-free water for by northern blot analysis, as previously described (98) with oligomers listed in **Table S2**.

### RNA sequencing

RNA samples from the gradient factions of naïve or ΦKZ-infected bacteria were diluted 1:10 with nuclease-free water and 10 μl of the dilution were mixed with 10 μl of 1:100 diluted ERCC RNA Spike-In Mix 1 (Thermo Fischer Scientific). This mix was used as an external RNA control for read normalization and to determine the dynamic range in sequencing, as suggested in best practice guidelines by the External RNA Controls Consortium (Baker et al. 2005). RNA-sequencing libraries were prepared by VERTIS Biotechnologie AG (Freisingen, Germany) as previously described (29). In essence, RNA samples were fragmented by ultrasound (four pulses of 30 s at 4°C) followed by 3’ adapter ligation. The 3’ adapter served as primer, synthesis of the first-strand cDNA was performed using M-MLV reverse transcriptase. After purification, 5’ Illumina TruSeq sequencing adapter was ligated to the end of the antisense cDNA. The resulting cDNA was PCR-amplified to about 10–20 ng/μl using a high-fidelity DNA polymerase and purified with the Agencourt AMPure XP Kit (Beckman Coulter Genomics). The cDNA samples were pooled based on the ratios according to the RNA concentrations of the input samples, and a size range of 200–550 bp was eluted from a preparative agarose gel. This size-selected cDNA pool was finally subjected to sequencing on an Illumina NextSeq 500 system using 75-nt single-end read length.

### Mass spectrometry

The nanoLC-MS/MS analysis was performed as previously described (29). Protein samples diluted in 1.25× protein loading buffer were homogenized by ultrasound [five cycles of 30 s on followed by 30 s off, high power at 4°C, Bioruptor Plus, Diagenode]. Insoluble material was removed by pelleting at full speed. A 20 μl sample of the cleared protein was mixed with 10 μl of UPS2 spike-in (Sigma-Aldrich) and diluted in 250 μl 1.25× protein loading buffer. The samples were subsequently reduced in 50 mM DTT for 10 min at 70°C and alkylated with 120 mM iodoacetamide for 20 min at room temperature in the dark. The proteins were precipitated in four volumes of acetone overnight at -20°C. Pellets were washed four times with ice-cold acetone and dissolved in 50 μl 8 M urea, 100 mM ammonium bicarbonate. Digestion of the proteins was performed by 0.25 μg Lys-C (Wako) for 2 h at 30°C, followed by dilution to 2 M urea by addition of 3 volumes 100 mM ammonium bicarbonate, pH 8 and overnight digestion with 0.25 μg trypsin at 37°C.

Peptides were purified through C-18 Stage Tips (99). Each Stage Tip was prepared with three disks of C-18 Empore SPE Disks (3M) in a 200 μl pipette tip. Peptides were eluted with 60% acetonitrile/0.3% formic acid, lyophilized in a laboratory freeze-dryer (Christ), and stored at -20°C. Prior to nanoLC-MS/MS, the peptides were dissolved in 2% acetonitrile/0.1% formic acid.

### NanoLC-MS/MS analysis

NanoLC-MS/MS analysis was performed as previously described (29) with an Orbitrap Fusion (Thermo Fisher Scientific) equipped with a PicoView Ion Source (New Objective) and coupled to an EASY-nLC 1000 (Thermo Fisher Scientific). Peptides were loaded on capillary columns (PicoFrit, 30 cm × 150 μm ID, New Objective) self-packed with ReproSil-Pur 120 C18-AQ, 1.9 μm (Dr. Maisch) and separated with a 140-min linear gradient from 3 to 40% acetonitrile and 0.1% formic acid at a flow rate of 500 nl/min.

Both MS and MS/MS scans were acquired in the Orbitrap analyser with a resolution of 60,000 for MS scans and 15,000 for MS/MS scans. HCD fragmentation with 35% normalized collision energy was applied. A Top Speed data-dependent MS/MS method with a fixed cycle time of 3 s was used. Dynamic exclusion was applied with a repeat count of 1 and an exclusion duration of 60 s. Precursors that were singly charged were excluded from selection. Minimum signal threshold for precursor selection was set to 50,000. Predictive AGC was used with a target value of 2 × 105 for MS scans and 5 × 104 for MS/MS scans. EASY-IC was used for internal calibration.

### Grad-seq RNA data analysis

Reads were mapped to the *Pseudomonas* and ΦKZ reference sequences (NC_002516 and NC_004629, respectively) using the READemption align function (READemption version 0.4.3, (100)). About ∼96% of reads were aligned, for details see the statistics files in the READemption analysis folder. Coverage wig-files were generated with the coverage function and read allocation to genomic features was quantified by the gene_quanti function (gff files are supplied in the READemption analysis folder, Zenodo 4024332). Annotation of 3’/5’UTRs was used from our parallel study and sRNAs from PseudoCAP (42), and manually annotated transcripts were added. Sequencing coverages were visualized in the Integrative Genomics Viewer (IGV, Broad Institute, (101)) based on uniquely mapped reads and normalized to the total number of aligned reads.

### Grad-seq MS data analysis

Raw MS data files were analysed with MaxQuant version 1.6.2.2 (102). Database search was performed with Andromeda, which is integrated in the utilized version of MaxQuant. The search was performed against the UniProt *Pseudomonas aeruginosa* UP000002438 (strain PAO1), the ΦKZ proteome UP000002098 (37) and a database containing the proteins of the UPS2 proteomic standard. In addition, a database containing common contaminants was used. The search was performed with tryptic cleavage specificity with three allowed missed cleavages. Protein identification was under control of the false-discovery rate (FDR, 1% on protein and peptide level). In addition to MaxQuant default settings, the search was additionally performed for the following variable modifications: Protein N-terminal acetylation, glutamine to pyro-glutamic acid formation (N-term. glutamine) and oxidation (methionine). Carbamidomethyl (cysteine) was set as fixed modification. For protein quantitation, the iBAQ intensities were used (103). Proteins that could not be distinguished by peptides were listed individually.

### Normalization and dose-response curves

Relative protein abundance in gradient fractions were calculated by iBAQ, as described previously (75). Relative protein abundance was estimated by correcting for alterations in digestion and C18 purification in fractions by normalization to human albumin (**Table S1**, norm. to spike-in, P02768ups|ALBU_HUMAN_UPS Serum albumin, chain 26-609, **Fig. S1A-C**) that was spiked-in as part of the UPS2 standard (Sigma-Aldrich). We selected albumin for normalization because of the highest number of recovered peptides (104). In bar-diagram representation, the abundances for each individual protein were transformed into distributions across the gradient by dividing the abundance in each fraction by the maximal abundance across the gradient (**Table S1**, norm. to max.). For visualization purposes of differential sedimentation representation, the y-axes of individual profiles were scaled to the largest value in both represented gradients. For RNAs basically the same approach was used with the ERCC-0130 spike in, as all spike-in RNAs behaved well correlated in sample-wise quantification. Based on the corrected quantification values, we setup a dose-response curve plotting the measured relative abundance vs. the input abundance (**Fig. S1D-F**). The linear range of recovery was determined for proteins being log_10_iBAQ >7.5 and for log_10_RNA levels >1.5. All proteins and RNAs that were recovered below the threshold were neglected representing 614 proteins (**Fig. S1B**) and 2,182 RNAs (**Fig. S1E**).

### Oligo probe labelling and northern blotting

Oligos were radioactively labelled at 5’-OH with [γ-^32^P]ATP by the T4-polynucleotide kinase, as reported previously (75). Northern probing was performed as previously reported (98).

### Data availability

MS data are accessible at the ProteomeXchange consortium (105) via the PRIDE partner repository (106) with the dataset identifier PXD021359. Raw data after MaxQuant and sequencing analysis are listed in **Table S1** in the tab ‘RAW’. Sequencing raw FASTQ and analysed WIG and TDF coverage files are accessible at Gene Expression Omnibus (GEO, (107)) with the accession no. GSE157708. The code for the Gradseq browser is deposited at Zenodo (DOI:10.5281/zenodo.3955585). The GradR browser is online accessible at https://helmholtz-hiri.de/en/datasets/gradseqpao1. READemption 0.4.5 is deposited at Zenodo 1134354 (DOI:10.5281/zenodo.1134354). The READemption analysis folder is deposited at Zenodo 4024332 (DOI:10.5281/zenodo. 4024332).

## ACKNOWLEDGEMENTS

We thank Barbara Plaschke for technical assistance, Stephanie Lamer, Andreas Schlosser, Jens Hör, Lars Barquist, Konrad Förstner and Silvia di Giorgio for helpful discussions about experiments and data analysis. We thank the Helmholtz Institute for RNA-based Infection Research (HIRI) who supported this work with a seed grant through funds from the Bavarian Ministry of Economic Affairs and Media, Energy and Technology (Grant allocation nos. 0703/68674/5/2017 and 0703/89374/3/2017). This work was also supported by a Gottfried Wilhelm Leibniz award (DFG Vo875/18). L.W. holds a predoctoral scholarship from FWO-fundamental research (11D8920N). This manuscript was supported by funding from the European Research Council (ERC) under the European Union’s Horizon 2020 research and innovation programme (Grant agreement No. 819800) awarded to R.L.

L.W., M.G., J.V. designed and performed experiments. M.G. analysed the Grad-seq data and carried out bioinformatics analyses and generated figures for the manuscript. M.G. and J.V. wrote the manuscript with input from all authors. J.V. supervised and guided the project and interpreted Grad-seq data. All authors approved the final version.

We declare no competing interests.

## SUPPLEMENTARY MATERIAL

**Figure S1:**
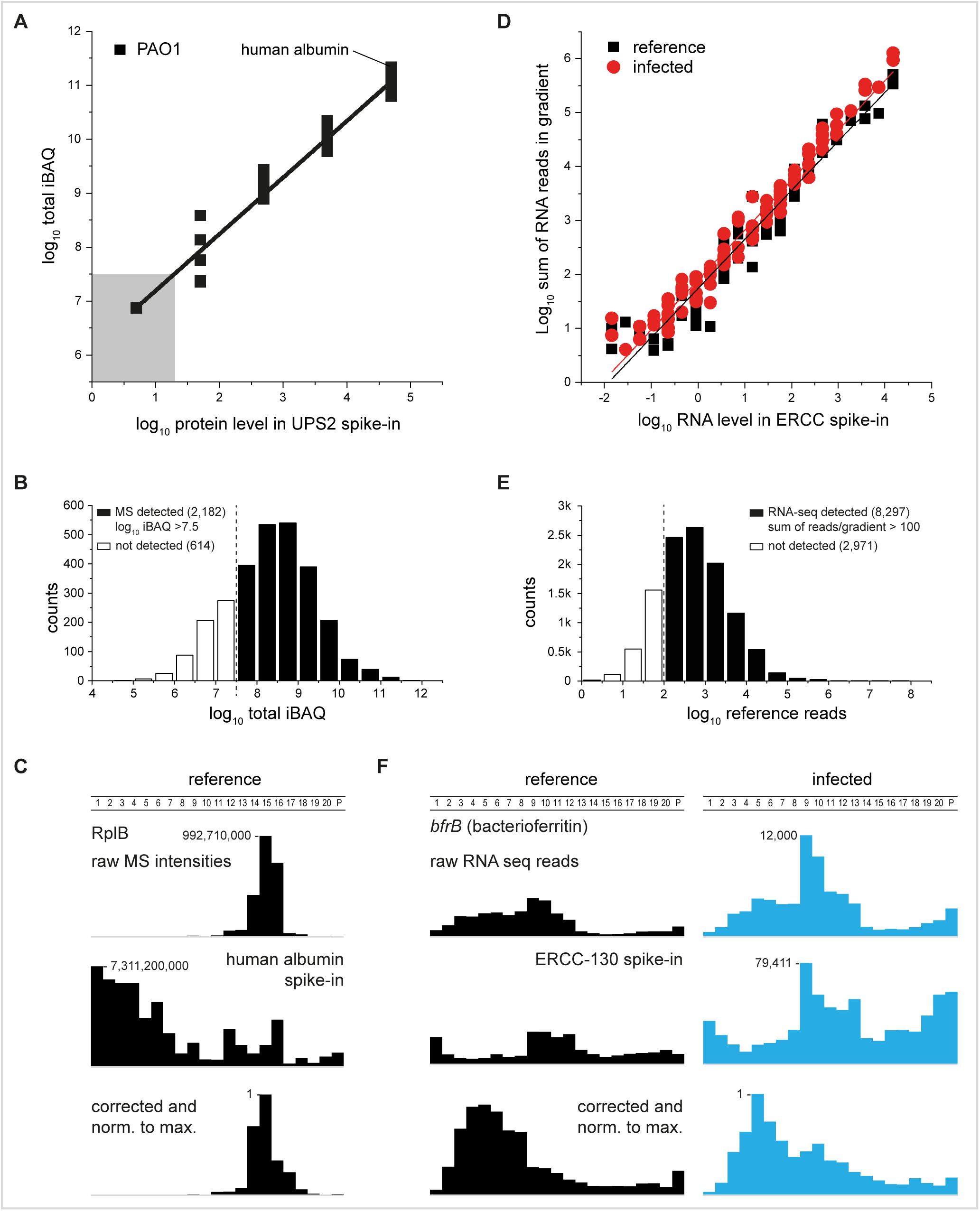
Analysis and normalization of Grad-seq. **(A)** Dose-response curve for the MS analysis, plotted as relative concentration in UPS2 spike-in vs. recovered MS counts. The linear dynamic range of detection is valid down to log_10_ total iBAQ of 7.5. **(B)** 2,182 proteins were selected above the threshold from (A). 614 proteins were discarded. **(C)** Normalization of protein levels by human albumin (UPS2) spike-in. **(D)** Dose-response curve for the RNA-seq analysis based on the ERCC spike-in. The linear dynamic range of detection was valid down to log_10_ sum of RNA reads of 1.5. **(E)** 8,297 annotations were detected above the threshold of >100 reads/gradient, 2,971 annotations were excluded. **(F)** Normalization of RNA levels by the ERCC-130 spike-in.

**Figure S2.**
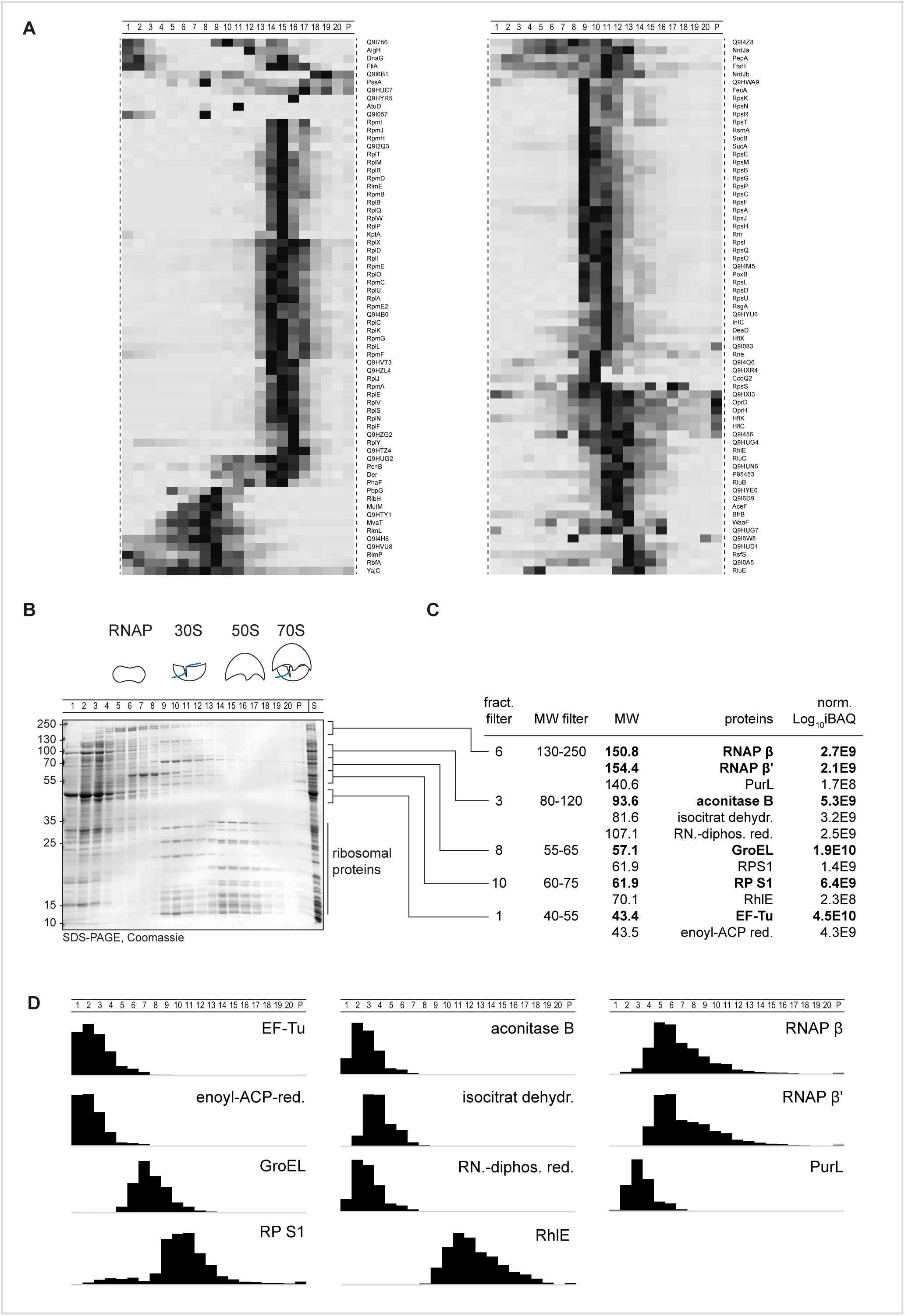
Sedimentation profiles of complexes that peak in fractions 8-20. **(A)** About 30 uncharacterized proteins sedimented in HMW fractions and are named here by UniProt identifier. **(B)** Selected abundant protein bands in the Grad-seq experiment were selected for identification. **(C)** Protein entries were filtered by selected fraction where the protein was mostly abundant in (B) and a range of molecular weight based on the molecular size marker. The resulting table was ranked by abundance resulting in candidates. The abundance (iBAQ) was evaluated in contrast to the second-best hit. **(D)** Digital sedimentation profiles were correlated with the sedimentation profiles in (B).

**Figure S3.**
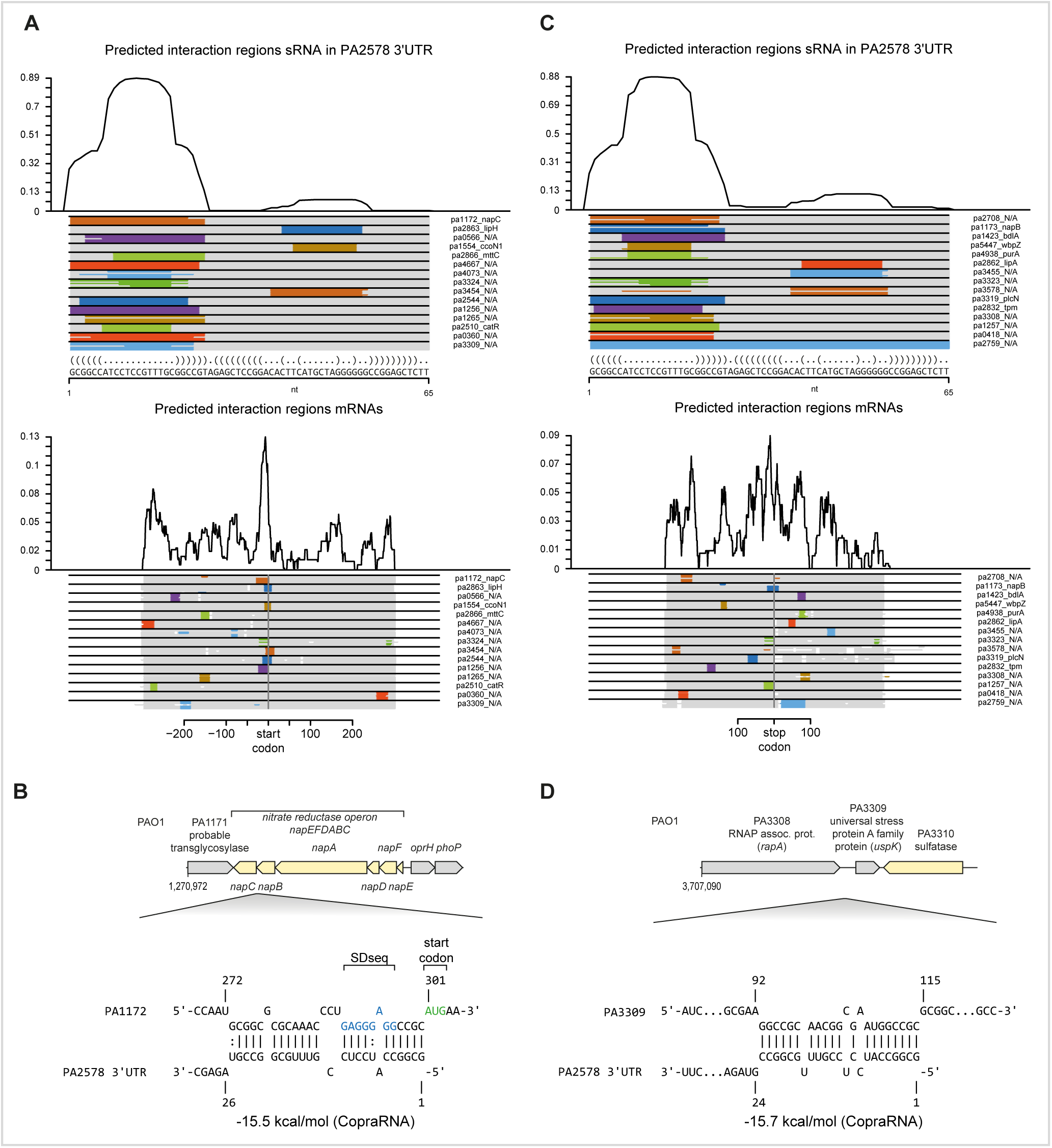
CopraRNA analysis of RNA-RNA interaction of the PA2758-3’UTR fragment. **(A)** Predicted interaction region around start codons were determined being in the first stem loop that may represent the seed sequence. **(B)** An example is the intergenic region between *napB* and *napC* where the fragment could pair with the Shine-Dalgarno-sequence (SDseq) and modulate translation initiation. **(C)** Predicted interaction regions around stop codons. **(D)** A second example covers an interaction in the 5’UTR of the universal stress protein A family protein *uspK*.

**Figure S4.**
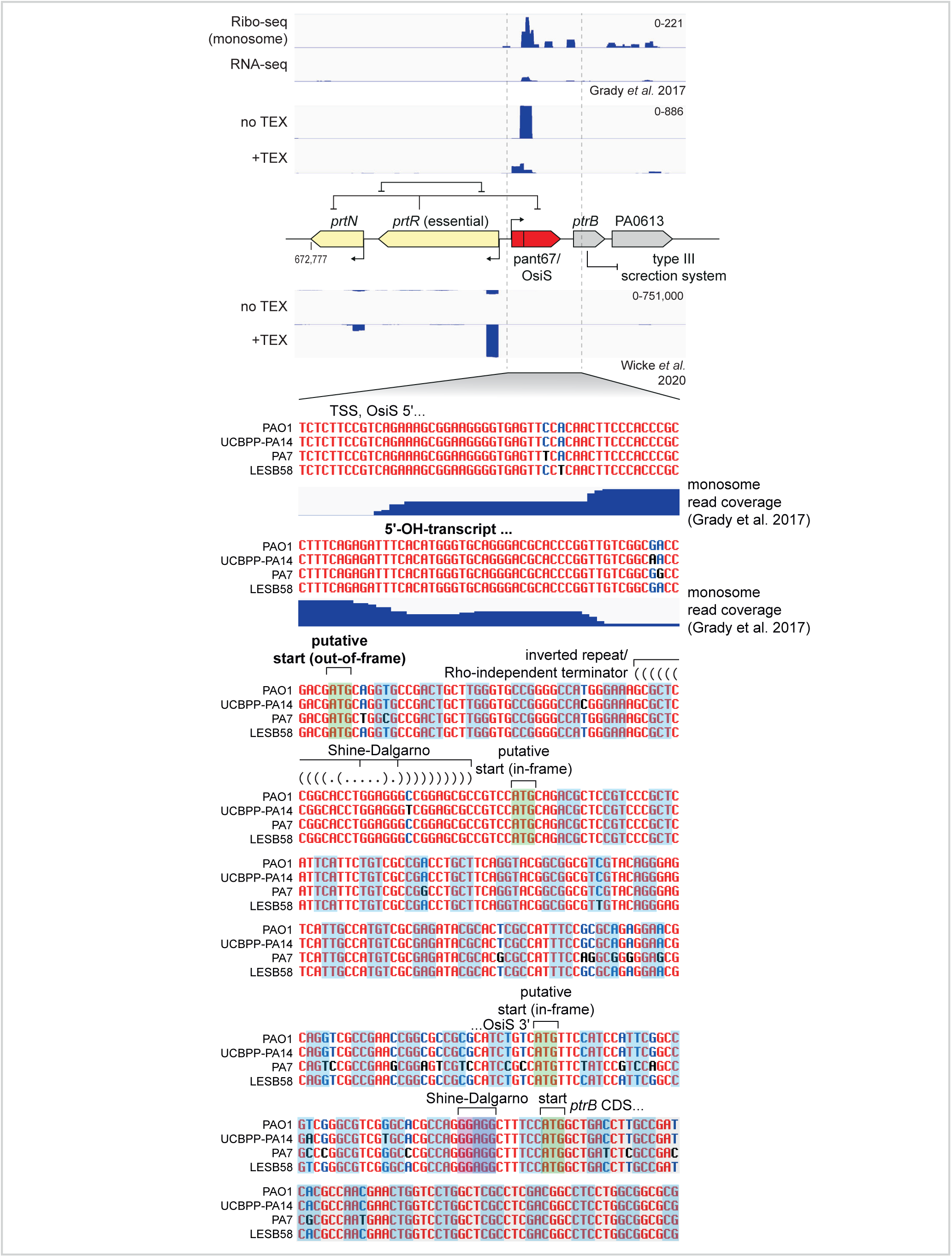
ncRNA OsiS may harbour a hidden small open reading frame that may drive sedimentation in 30S fractions. TEX treatment indicates a TEX unaffected 5’-end downstream of the annotated TSS (51). Sequencing of RNA fragments recovered from monosomes (Ribo-seq, (66), re-analysed with HRIBO (https://github.com/RickGelhausen/HRIBO)) supports the possibility of a short translated fragment starting at a putative start codon that is out-of-frame with *ptrB*. In addition, an inverted repeat in-between may serve as a Rho-independent terminator because no stop-codon is reached in the reading frame.

**Figure S5.**
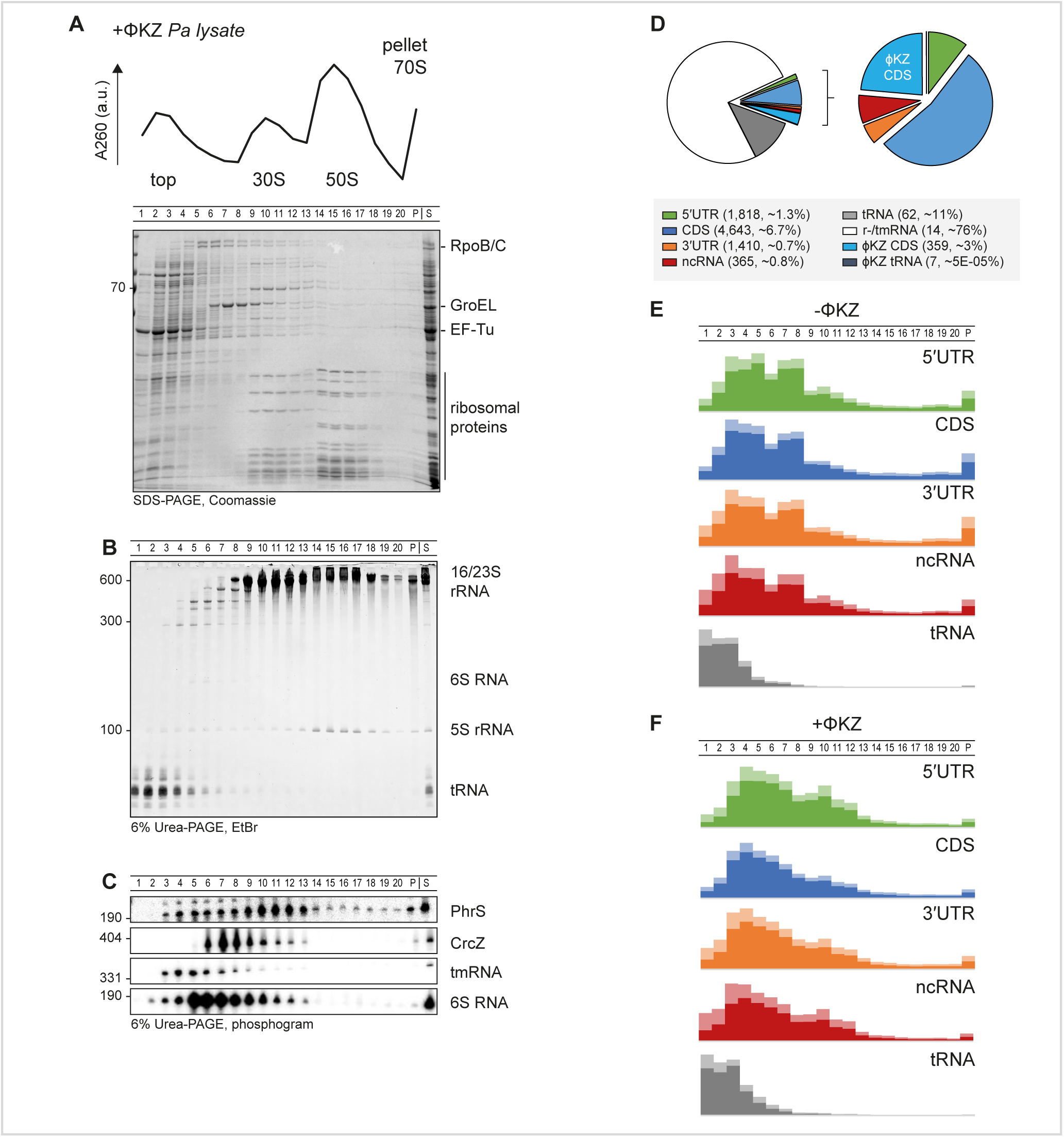
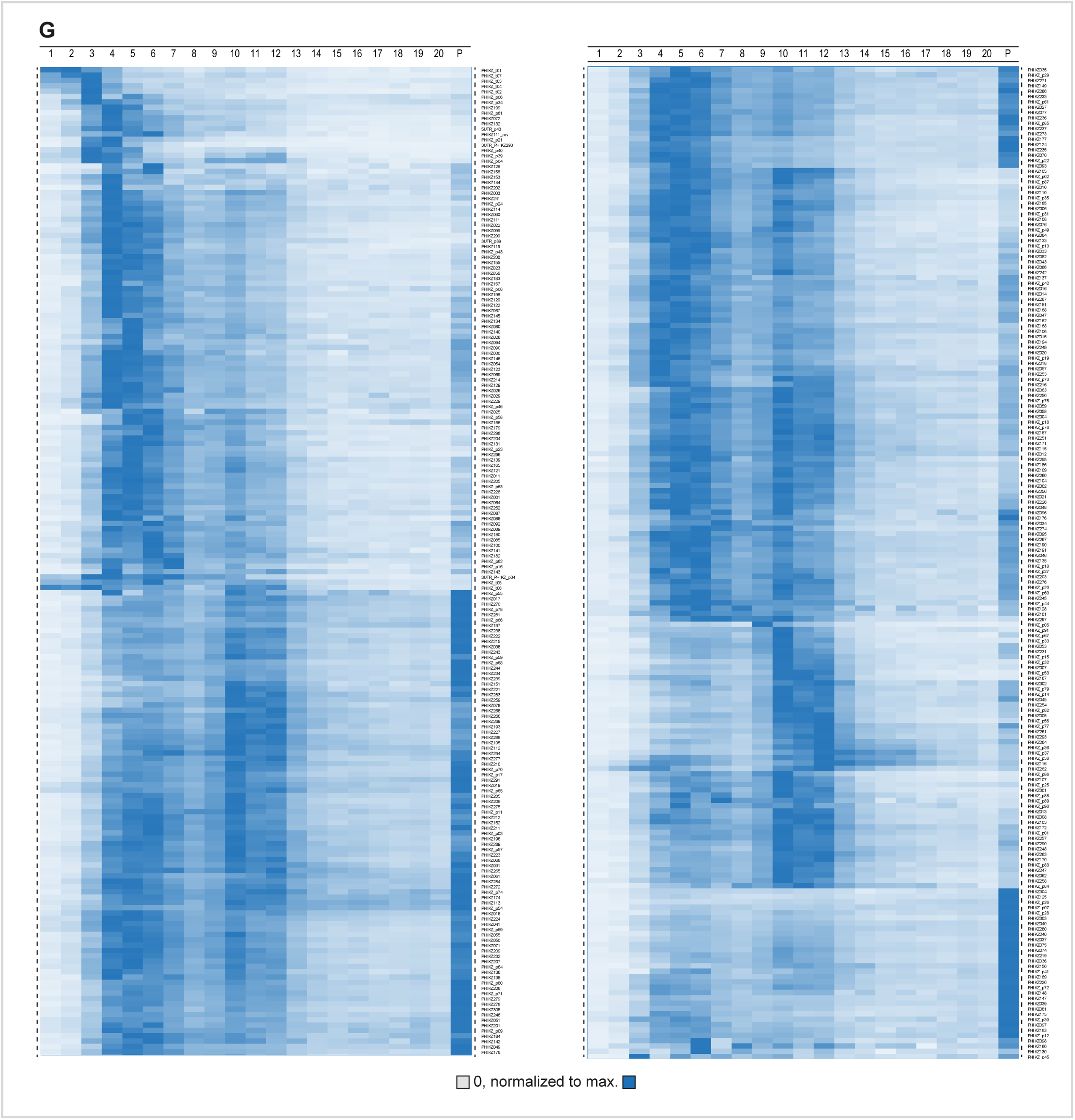
Gradient fractionation of ΦKZ-infected *Pseudomonas* at early infection time point. (A) The absorption profile and apparent proteome is not affected upon infection. **(B)** The global picture of the RNome is consistent with Fig. 1D and not altered upon infection. **(C)** Specific RNAs were probed by northern blotting and resulted in consistent sedimentation behaviour with non-infected cells. **(D)** Phage transcripts represented about 3% of all reads. Host CDS transcripts remained present at 7% - relative representation. **(E)** In non-infected cells, the sedimentation profile of CDS, UTRs, and ncRNAs was strongly balanced on the side of RNAP fractions and marginally on ribosomal fractions. tRNAs sedimented exclusively in LMW fractions. **(F)** In infected cells, the sedimentation profiles of the host transcripts were only marginally affected and shifted towards LMW fractions. **(G)** Phage transcripts sedimented strongly in ribosomal fractions.

**Figure S6.**
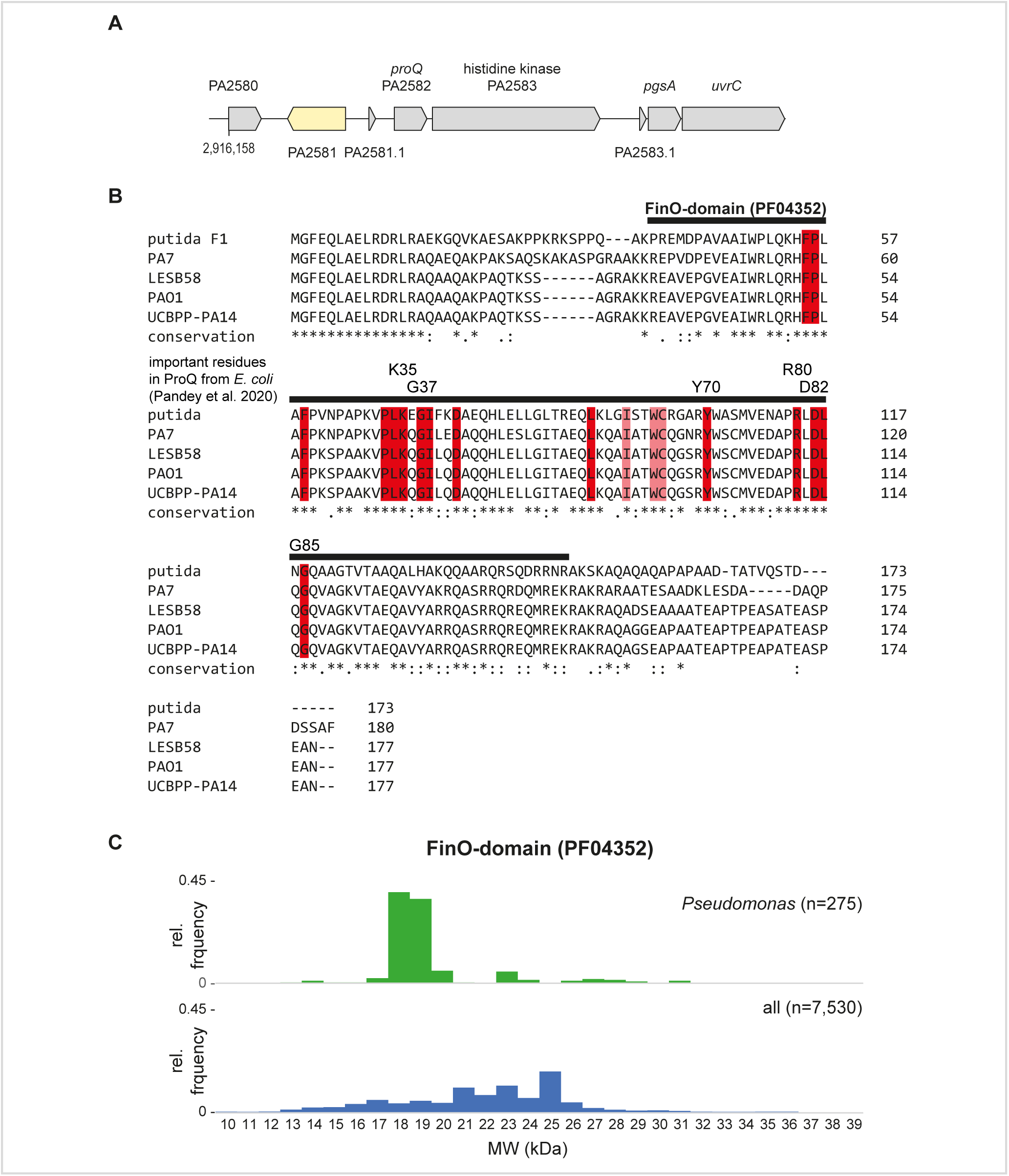
ProQ in *Pseudomonas* harbours important residues of a FinO-domain. **(A)** Genomic locus of ProQ in PAO1. **(B)** Conserved residues in a FinO-domain (PF04352) are represented in red. Important residues in ProQ for RNA-binding (76) are indicated with corresponding residues in ProQ from *E. coli.* **(C)** ProQ proteins in *Pseudomonas* represent the condensed representatives in the diverse class of FinO-domain proteins that can have C- or N-terminally associated domains.

**Figure S7.**
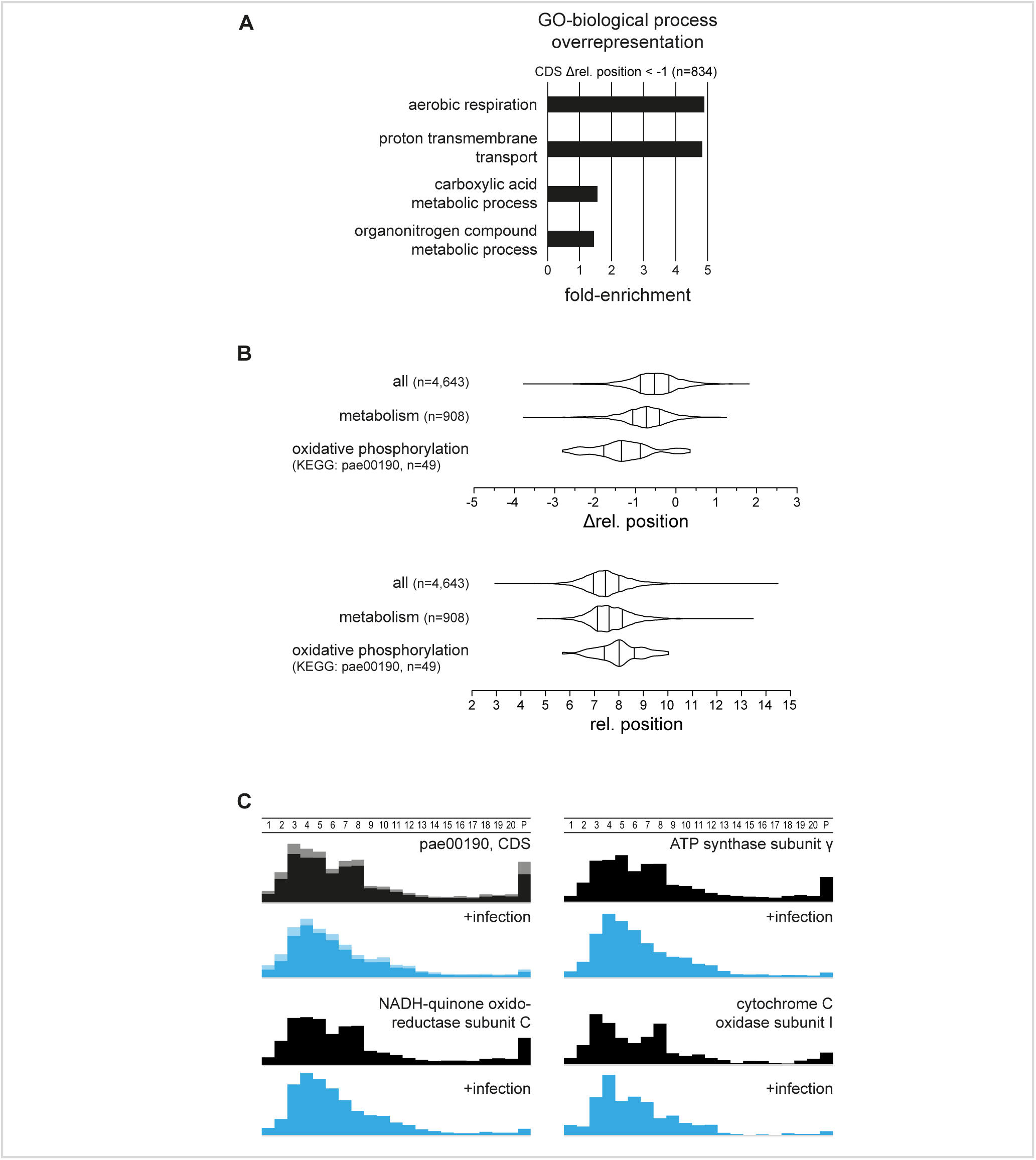
Transcripts for oxidative phosphorylation are shifted most strongly towards LMW fractions. **(A)** Overrepresentation analysis for biological processes revealed that 5’UTR transcripts that are shifted by >0.5 rel. fractions are enriched for transcription factors. 3’UTR transcripts that are shifted by more than 1 rel. fraction towards the top are enriched for factors of aerobic respiration and proton transmembrane transport. **(B)** 49 transcripts encoding oxidative phosphorylation factors are significantly more shifted towards the top compared to transcripts coding for metabolic factors. At the same time the relative position in the reference condition was for these transcripts significantly higher and matches the trend in (A). **(C)** Averaged sedimentation profile of the oxidative phosphorylation pathway pae00190 CDS, and selected examples coding for ATP synthase, NADH-oxidoreductase, and cytochrome C oxidase.

**Figure S8.**
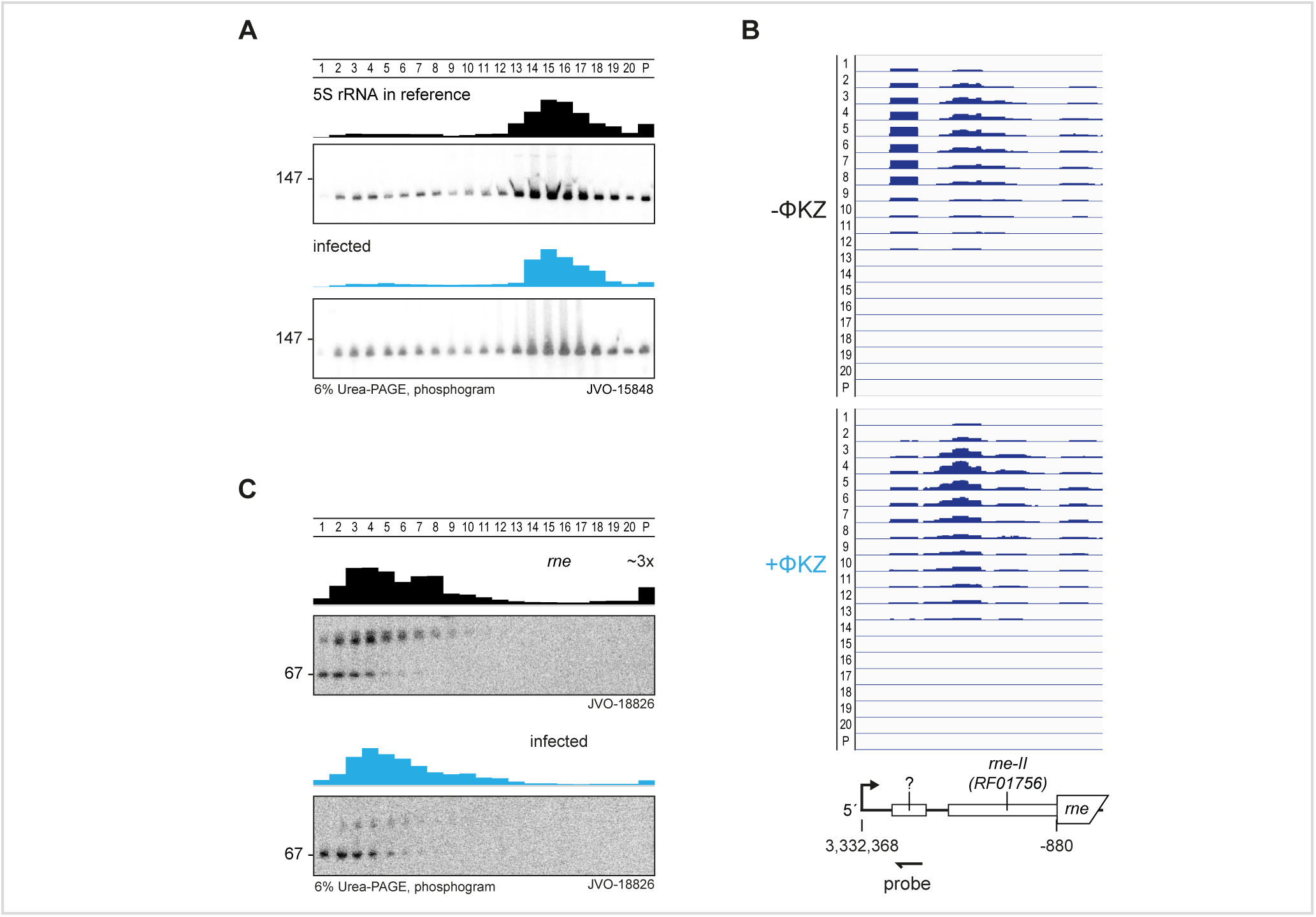
5S rRNA indicates processing effects and its major RNase E is altered in the 5’UTR. **(A)** 5S rRNA sedimentation was not affected upon infection, but it become smeary in northern probing that may indicate processing defects that could lead to diminished reads in sequencing. **(B)** The read-coverage of the 5’UTR of *rne* was altered upon infection. **(C)** The 5’UTR appeared in two fragments of which the larger one was allocated to RNAP fractions and strongly diminished upon ΦKZ infection.

Table S1. Grad-MS & -seq data (xls file)

Table S2. STAR methods (xls file)

